# *In vivo* measurements of receptor tyrosine kinase activity reveal feedback regulation of a developmental gradient

**DOI:** 10.1101/2025.01.06.631605

**Authors:** Emily K. Ho, Rebecca P. Kim-Yip, Alison G. Simpkins, Payam E. Farahani, Harrison R. Oatman, Eszter Posfai, Stanislav Y. Shvartsman, Jared E. Toettcher

**Author notes:** Corresponding Authors: Emily Kolenbrander Ho Lewis Thomas Laboratory Room 141 Washington Road Princeton, NJ 08544 Jared Toettcher Lewis Thomas Laboratory Room 140 Washington Road Princeton, NJ 08544.

## Abstract

A lack of tools for detecting receptor activity *in vivo* has limited our ability to fully explore receptor-level control of developmental patterning. Here, we extend a new class of biosensors for receptor tyrosine kinase (RTK) activity, the pYtag system, to visualize endogenous RTK activity in *Drosophila*. We build biosensors for three *Drosophila* RTKs that function across developmental stages and tissues. By characterizing Torso::pYtag during terminal patterning in the early embryo, we find that Torso activity differs from downstream ERK activity in two surprising ways: Torso activity is narrowly restricted to the poles but produces a broader gradient of ERK, and Torso activity decreases over developmental time while ERK activity is sustained. This decrease in Torso activity is driven by ERK pathway-dependent negative feedback. Our results suggest an updated model of terminal patterning where a narrow domain of Torso activity, tuned in amplitude by negative feedback, locally activates signaling effectors which diffuse through the syncytial embryo to form the ERK gradient. Altogether, this work highlights the usefulness of pYtags for investigating receptor-level regulation of developmental patterning.

## Introduction

Tissues are patterned by ligand-receptor signaling. In the classic model of developmental morphogen patterning, a local source of ligand patterns distinct domains of cell identity by diffusing across a field of cells and activating receptors in a concentration-dependent manner (Briscoe and Small, 2015; Wolpert, 1969). These receptors then activate conserved signaling cascades that induce different target genes at different thresholds, thereby converting the positional information encoded in ligand-receptor interactions into spatial patterns of gene expression.

Recent advances in imaging and biosensor development have deepened our understanding of the cellular processes that produce these patterns. High-resolution detection of ligand trafficking, including single-molecule imaging, has shown that ligands do not always move via simple diffusion but associate significantly with cell membranes (Hall et al., 2021; Kuhn et al., 2022; Schlissel et al., 2024). Live biosensors of signal transduction, such as kinase translocation reporters, have uncovered heterogeneous and dynamic patterns of pathway activity controlling essential fate decisions (De La Cova et al., 2017; Regot et al., 2014; Wilcockson et al., 2023). And tools for measuring gene expression, such as the MS2-MCP system, have revealed how target genes tune their transcriptional kinetics in response to inputs of different strengths rather than simply reaching a binary expression threshold (Carmon et al., 2021; Dubrulle et al., 2015; Garcia et al., 2013; Ho et al., 2023). Thus, developmental patterns emerge from complex signal processing at multiple pathway nodes.

Significant signal processing can also occur at the level of the receptors themselves. For example, the location and level of receptor expression can alter ligand distribution (De Vreede et al., 2022; Eldar et al., 2003; Kicheva and Briscoe, 2023; Zhu et al., 2020). Accumulating evidence suggests that receptor *activity* is also highly regulated and not simply a function of ligand concentration. For example, the activity state of receptor tyrosine kinases (RTKs) is known to be influenced by mechanisms such as feedback regulation of receptor phosphorylation (Neben et al., 2019), negative cooperativity in receptor dimer formation (Alvarado et al., 2010; Jenni et al., 2015), and ligands with different binding affinities (Deguchi et al., 2024; Freed et al., 2017). However, the consequences of these mechanisms for tissue patterning have largely remained unexplored due to a lack of tools for monitoring receptor activity *in vivo*. Further, the RTK-associated Ras/ERK signaling pathway not only forms long-range gradients but also pulses and traveling waves (Aoki et al., 2017; De Simone et al., 2021; Lim et al., 2015; Valon et al., 2021). How receptor activity varies across space and time to produce these patterns remains unclear. Therefore, new tools for directly detecting receptor activity *in vivo* are greatly needed, and RTK biosensors would be of particular interest in many developmental contexts.

Our recent investigations into a canonical RTK patterning event in the *Drosophila melanogaster* embryo spurred our efforts to build new tools for monitoring RTK activity *in vivo*. In the first 3 hours of development, the maternal terminal patterning system defines the poles, or termini, of the syncytial embryo as the future head and tail (Smits and Shvartsman, 2020). This patterning system involves a maternally deposited RTK, Torso, and its ligand Trunk (Casali and Casanova, 2001; Casanova and Struhl, 1989; Casanova et al., 1995). Torso and Trunk are present throughout the embryo, but Trunk is only processed into its active form at the poles, resulting in localized Torso binding, activation of the Ras/ERK cascade, and gene expression patterning (Furriols et al., 1996; Ghiglione et al., 1999; Greenwood and Struhl, 1997; Lu et al., 1993; Sprenger et al., 1993). Although the ERK gradient is well characterized, the pattern of Torso activity has never been directly visualized in the embryo (Coppey et al., 2008). Through optogenetic approaches, we recently discovered that a simple, all-or-none pattern of Ras activation at the poles can produce gradients of intracellular signaling and domains of gene expression that are sufficient for normal terminal patterning and subsequent development (Ho et al., 2023; Johnson et al., 2020). This surprising result suggested that diffusion of pathway effectors in the syncytial embryo can create a discrepancy between the spatial domain of receptor-level activity and the resulting pattern of downstream signaling. Thus, terminal patterning by Torso is an ideal model to investigate RTK activity, because receptor activity, in principle, might be quite different from the pattern of downstream ERK activity.

Here, we address the lack of tools for visualizing endogenous RTK activity by extending a new class of biosensors for RTK activity, the pYtag system, to *Drosophila*. We show that pYtags enable live visualization of *Drosophila* RTK activity across multiple receptors, developmental stages, and tissues. We then use the Torso::pYtag system to characterize Torso activity during terminal patterning for the first time. We find that Torso activity differs from ERK activity in two important ways: Torso activity decreases and recedes to the pole during a developmental time window when terminal ERK activity is increasing, and ERK activity extends beyond the predominant domain of active Torso. Drawing upon these observations, we show that Torso activity is regulated by Ras/ERK-pathway dependent negative feedback, suggesting that a feedback loop between Torso and ERK shapes the terminal pattern. We propose that blurring of a local activity domain into a longer range ERK gradient decouples regulation of signal amplitude from gradient formation. Together, this work provides a new model for terminal patterning and highlights the usefulness of pYtag RTK biosensors.

## Results

### Design of endogenous pYtag biosensors for *Drosophila* RTKs

RTKs such as Torso transduce extracellular ligand binding through phosphorylation of tyrosines on their receptor tails. These phosphorylated tyrosines (pYs) bind to Src homology 2 (SH2)-domain containing signaling effectors, triggering downstream signaling (Lemmon and Schlessinger, 2010). Prior studies have deployed SH2-domain-based RTK biosensors to detect changes in receptor phosphorylation state, but these biosensors typically lack specificity due to promiscuous phosphotyrosine-SH2 interactions, and competition for receptor binding between the biosensor and endogenous SH2-domain containing proteins can disrupt signaling (Ladbury and Arold, 2000; Tiruthani et al., 2019; Wintgens et al., 2019).

We recently addressed both limitations with pYtags, new RTK biosensors that take advantage of an exceptionally specific phosphotyrosine-binding domain interaction found in immune cells (Farahani et al., 2023). T-cell receptors harbor immunoreceptor tyrosine-based activation motifs (ITAMs) containing pairs of tyrosines that, when phosphorylated, selectively recruit the tandem SH2 domain of ZAP70 (ZtSH2) (Isakov et al., 1995; Wange et al., 1993). This ITAM-ZtSH2 interaction is 100-fold more specific than interactions between the ITAM and a single ZAP70 SH2 domain and, given its exclusive expression in immune cells, provides an orthogonal pair with which to build a biosensor (Isakov et al., 1995). We showed in mammalian cells that appending 3 copies of the CD3ε ITAM sequence (3x CD3ε) to the C-terminus of an RTK creates a selective binding site for a fluorescently-tagged ZtSH2 (**Figure 1A**) (Farahani et al., 2023). In this manner, receptor phosphorylation can be monitored by the cytoplasmic-to-membrane translocation of the fluorescent ZtSH2 biosensor in live cells. In principle, pYtags are applicable to any RTK of interest to yield a specific, dynamic readout of receptor activity, but can they provide a readout of RTK activity *in vivo*?

**Figure 1:**
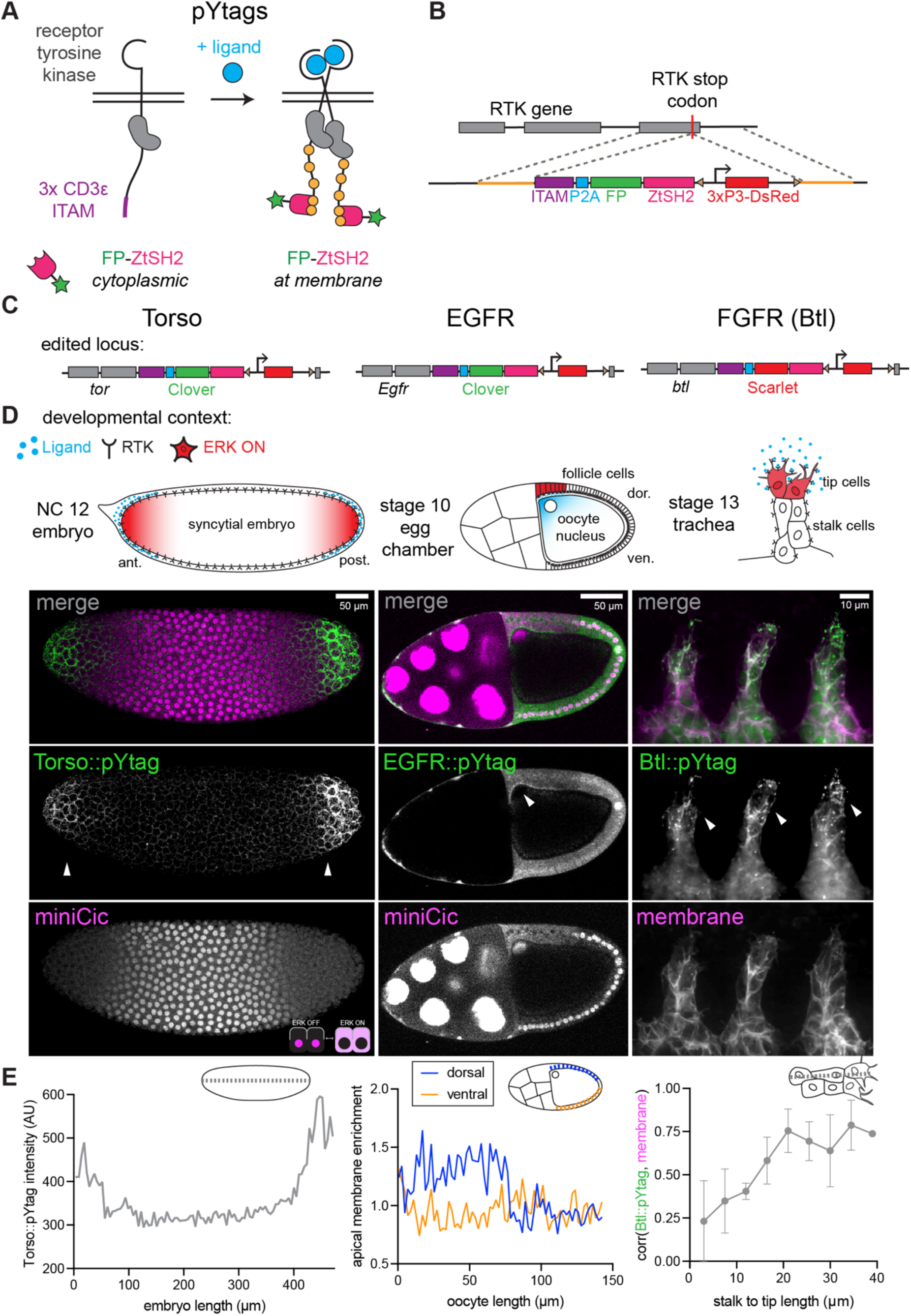
pYtags are endogenous biosensors of RTK activity. (A) Design of pYtag biosensors. An ITAM sequence (103 amino acids) is appended to the C-terminal tail of an RTK. Upon ligand binding, phosphorylation of tyrosines in the ITAM leads to specific membrane recruitment of a fluorescently-tagged tandem SH2 protein. FP = fluorescent protein (B) CRISPR/Cas9 insertion of the pYtag into *Drosophila* RTK genes. The 5.5 kb homology construct contains the pYtag cassette, a 3xP3-DsRed flanked by piggyBac sites (triangles) for detecting the insertion, and 1 kb homology arms (orange). (C) Edited loci for three RTKs: *Torso::pYtag*, *EGFR::pYtag*, and *Btl::pYtag*. (D) Imaging of each pYtag in a relevant tissue. Arrowheads denote regions of pYtag membrane recruitment and active RTK signaling. Left: Torso::pYtag in a nuclear cycle (NC) 12 embryo imaged together with miniCic, which translocates from the nucleus to the cytoplasm when ERK is active (see inset diagram), shows Torso activity at the poles. Center: EGFR::pYtag in a stage 10 egg chamber imaged together with miniCic shows EGFR activity in the dorsal follicle cells next to the oocyte nucleus. Right: Btl::pYtag in a stage 13 trachea imaged with a CD4::mIFP membrane marker shows Btl activity in the leading tip cells. (E) Quantification of each pYtag biosensor in the displayed image. Left: Intensity of Torso::pYtag across the anterior-posterior axis measured by line scan. Center: Apical membrane enrichment of EGFR::pYtag in dorsal and ventral follicle cells, calculated from membrane and cytoplasmic intensity measurements along the oocyte length. Right: Pearson’s correlation between Btl::pYtag and membrane intensity in 9 evenly-spaced ROIs along the tracheal branch from stalk to tip. Points show the Mean ± s.e.m of the three branches in the image.

To test if pYtags could be applied to detect Torso RTK activity in the *Drosophila* embryo, we designed a CRISPR/Cas9 strategy to endogenously tag the C-terminus of *Drosophila* RTKs with the pYtag cassette. We opted for endogenous tagging because RTKs are generally expressed at low levels and overexpression can result in ligand-independent activation (Du and Lovly, 2018). The homologous recombination cassette contains the 3x CD3ε ITAM sequence, a P2A self-cleaving peptide sequence, and a fluorescently tagged ZtSH2 (**Figure 1B**). In this design, the RTK of interest is endogenously tagged with the ITAM, and the tagged ZtSH2 is expressed in a 1:1 ratio with the RTK. The ratio of the two pYtag components (RTK-ITAM and ZtSH2) is important to consider because it impacts the amplitude of the biosensor’s response (Farahani et al., 2023). The cassette also contains a 3xP3-DsRed for detecting the insertion (Gratz et al., 2015).

Using this strategy, we successfully obtained *Torso::pYtag* flies (**Figure 1C** - left). These flies were homozygous viable and fertile, indicating that the pYtag does not substantially alter Torso function. To examine the effect of the pYtag cassette on signaling more carefully, we stained *Torso::pYtag* embryos for two ERK target genes *tailless (tll)* and *huckebein (hkb)*, whose boundaries of expression are set by the ERK gradient (Ghiglione et al., 1999; Greenwood and Struhl, 1997). Compared to wild-type flies, *Torso::pYtag* embryos showed *tll* and *hkb* boundaries shifted towards the center of the embryo, revealing a slight increase in terminal signaling that is nevertheless compatible with successful development (**Figure S1A**). To test whether this increase in signaling is due to the pYtag or the 3xP3-DsRed transgene in the homologous repair construct, we removed the 3xP3-DsRed transgene by PiggyBac-mediated transposition to make *Torso::pYtag[-DsRed]* embryos. These embryos showed a partial rescue of the wild type *tll* and *hkb* boundaries, suggesting that the DsRed transgene does influence Torso activity, likely by altering its expression levels (**Figure S1A**). The residual phenotype in *Torso::pYtag[-DsRed]* embryos may be due to the pYtag itself or disruption to Torso by a C-terminal tag. Given the usefulness of maintaining the DsRed for subsequent genetic crosses, we use the original, DsRed-containing *Torso::pYtag* line for all subsequent experiments and confirm key results in *Torso::pYtag[-DsRed]* embryos.

To functionally test the Torso::pYtag biosensor, we live imaged homozygous *Torso::pYtag* embryos at 1-3 h post egg lay during Torso-mediated ERK activation at the embryonic poles (**Figure 1D** - left). We observed bright membrane enrichment of Clover-ZtSH2 at the poles but not in the center of the embryo, indicating that there are active Torso receptors with phosphorylated ITAMs at the poles (**Figure 1D** - left). We combined Torso::pYtag with the ERK activity kinase translocation reporter, miniCic, which is nuclear when ERK is off and cytoplasmic when ERK is on, to show that ERK is also active at the poles (Moreno et al., 2019).

Although the pYtag is designed to translocate between the cytosol and membrane, we detected a large change in membrane intensity at the termini but a uniformly low cytosolic background at all embryonic positions. This observation is likely due to the large cytosolic volume in the syncytial embryo, resulting in low cytosolic biosensor concentrations at all positions. Therefore, to quantify membrane recruitment of Clover-ZtSH2, we simply measured intensity across the anterior-posterior axis. These measurements of Clover-ZtSH2 intensity, which we refer to as “Torso::pYtag intensity”, revealed two peaks of Torso activity at the poles (**Figure 1E** – left). We observed similar peaks of Torso activity in *Torso::pYtag[-DsRed]* embryos (**Figure S1B)**. We also observed that the germ cells at the posterior pole (“pole cells”) lacked both Torso and ERK activity, consistent with previously-described degradation of Torso in these cells (**Figure S1C**) (Pae et al., 2017).

### Application of pYtags to other *Drosophila* RTKs

RTKs are a large family of receptors: humans have 58 RTKs and *Drosophila* have 20 (Lemmon and Schlessinger, 2010; Sopko and Perrimon, 2013). pYtags are a powerful class of biosensors because they are modular and can be appended to any RTK in principle without the need for receptor-specific design changes. We thus asked whether we could use the same strategy to detect the activity of other *Drosophila* RTKs *in vivo*. We inserted the pYtag into EGFR (*Egfr*), and the FGFR homolog Breathless (*btl*). Both lines were also homozygous-viable and fertile.

We assessed EGFR::pYtag in live stage 10 egg chambers, where the EGFR ligand Gurken is secreted near the oocyte nucleus to the overlying follicle cells to determine the oocyte’s dorsal side (**Figure 1D** – center) (Nilson and Schüpbach, 1998). We observed recruitment of Clover-ZtSH2 from the cytosol to the apical membrane of the dorsal follicle cells near the oocyte nucleus (**Figure 1D** – center) (Chang et al., 2008; Revaitis et al., 2020). Cytoplasmic miniCic in these follicle cells confirmed that ERK is active in the same follicle cells with active EGFR. Quantification of the apical enrichment of EGFR::pYtag showed that, as expected, EGFR is active in the dorsal follicle cells near the oocyte nucleus but not in the ventral follicle cells (**Figure 1E** – center).

We assessed Btl::pYtag during tracheal development. Btl guides migration of tracheal branches towards Branchless ligand (**Figure 1D** – right) (Sutherland et al., 1996). ERK signaling is active in the two leader “tip” cells of each branch but not in the trailing “stalk” cells (Hayashi and Kondo, 2018). Using Btl::pYtag, we observed recruitment of Scarlet-ZtSH2 to the membranes of tip cells and cytoplasmic localization in the stalk cells (**Figure 1D** – right). We assessed colocalization of Scarlet-ZtSH2 with membrane and found the two signals were highly correlated in the tip of the branch but not in the stalk (**Figure 1E** – right). We also noted an abundance of ZtSH2+ puncta in the Btl::pYtag trachea which we predict are signaling active endosomes (Haugh et al., 1999). Altogether, these three different *Drosophila* pYtags show that pYtags are broadly useful *in vivo* RTK biosensors across developmental stages and tissue types.

### Validating the efficacy and specificity of Torso::pYtag

The exquisite specificity of the pYtag biosensor derives from its orthogonality to endogenous SH2-pTyr interactions. However, *Drosophila* embryos express the ITAM-containing protein Downstream of kinase (Dok) and its binding partner Shark, the homolog of human ZAP70, which contains a tandem SH2 domain (Biswas et al., 2006; Fernandez et al., 2000). We thus asked if any of the observed membrane recruitment of Clover-ZtSH2 occurs because of interactions with a protein other than Torso. We noticed that even in ERK-OFF regions in the center of Torso::pYtag embryos, there was low-level membrane localization of Clover-ZtSH2 (**Figure 2A**), and this signal increased over NC11-14 (**Figure S2A, S2B**). This signal is ligand-independent, as *tsl^4^* embryos lacking *torso-like (tsl)*, the activator of the Trunk ligand, show no significant difference in Clover- ZtSH2 intensity in the center of the embryo (**Figure 2A, S2A, S2B**) (Savant-Bhonsale and Montell, 1993). Is this localization the result of non-specific recruitment of Clover-ZtSH2 to a protein other than Torso? We designed an “ITAM-less” version of Torso::pYtag that expresses only the Clover- ZtSH2 component and observed no membrane recruitment in the ITAM-less embryos, even in the center of the embryo (**Figure 2B**). Therefore, Clover-ZtSH2 is specifically recruited to the ITAM- tagged Torso, but there is a global baseline level of ligand-independent phosphorylation of Torso’s ITAM that is not indicative of Torso signaling. In all subsequent measures of Torso::pYtag activity, we account for this baseline activity by subtracting the Clover-ZtSH2 intensity in the center of the embryo.

**Figure 2:**
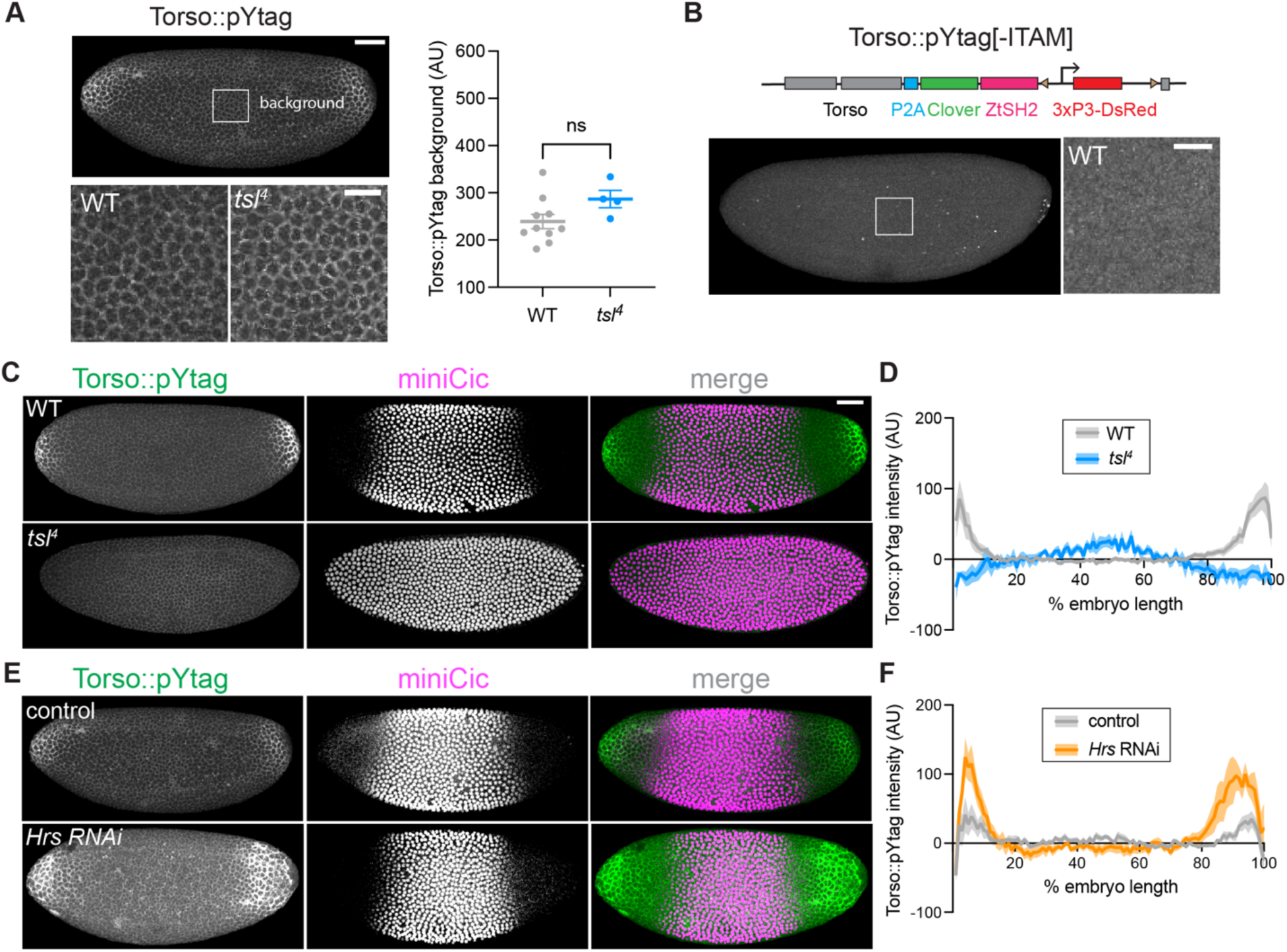
Torso::pYtag specifically detects Torso activity with relevant dynamic range. (A) Top image of a Torso::pYtag embryo highlights the inset region in the center of the embryo where Torso is inactive but background recruitment of Clover-ZtSH2 is observed. Bottom images show insets of WT and *tsl^4^* embryos in NC13 in this region. Graph shows no significant difference in the total Clover-ZtSH2 intensity of the inset region between genotypes. Mean ± s.e.m., n = 10, 4 embryos. Unpaired t test. (B) Without the ITAM sequence, Torso::pYtag[-ITAM] embryos show no membrane localization of Clover-ZtSH2. Inset zooms into the center of embryo in NC13. Diagram shows design of the Torso::pYtag[-ITAM]. (C, E) Comparison of Torso::pYtag and miniCic in WT versus *tsl^4^* mutant embryos and between control and *Hrs* RNAi embryos, all in NC13. For the miniCic channel only, the image look-up table differs to account for differences in miniCic copy number between conditions. (D, F) Quantification of Torso::pYtag intensity across the embryo length for each condition in NC13. Line shows mean ± s.e.m. of n embryos, n = 10 (WT), 4 (*tsl*), 8 (control), 9 (*Hrs* RNAi). Scale bars: 50 μm for whole embryo, 20 μm for insets.

We next sought to validate Torso::pYtag by showing that known perturbations to Torso activity result in expected changes to biosensor activity. We compared live, wild-type Torso::pYtag embryos with *tsl^4^* Torso::pYtag embryos in which Torso should be uniformly inactive across the embryo (Savant-Bhonsale and Montell, 1993). Torso-like (Tsl) is necessary for Trunk activity, and *tsl^4^* embryos have no terminal ERK activity as shown by nuclear miniCic throughout the embryo (**Figure 2C, S2D, S2E**). In *tsl^4^* embryos, we observed uniformly low Clover-ZtSH2 across the embryo with no enrichment at the poles, consistent with a lack of active Torso (**Figure 2C**). Quantification of Torso::pYtag intensity across the anterior-posterior axis, revealed that compared to WT embryos, which have two distinct peaks of Torso activation at the poles, *tsl^4^* embryos lack Torso activation (**Figure 2D**). Similar results were obtained for *trunk* mutant embryos (**Figure S2G, S2H**).

We also asked if perturbations that increase terminal signaling show increases in Torso::pYtag activity. Hepatocyte growth factor regulated tyrosine kinase substrate (Hrs) is an ESCRT-0 complex component necessary for degradation of active RTKs (Jékely and Rørth, 2003; Lloyd et al., 2002), and embryos lacking Hrs have increased terminal ERK signaling (Lloyd et al., 2002). Using RNAi to deplete maternal Hrs, we observed a significantly smaller domain of nuclear miniCic consistent with increased ERK signaling, although the effect was variable (**Figure 2E, S2F**). We also observed an expanded domain of Torso::pYtag membrane enrichment in *Hrs* RNAi embryos (**Figure 2E**). Quantification of Torso::pYtag intensity showed a larger domain of active Torso in *Hrs* RNAi embryos compared to control embryos expressing the maternal Gal4 but no RNAi (**Figure 2F**). We also note that *Hrs* RNAi embryos had a significantly increased background level of Torso::pYtag intensity compared to the RNAi controls, suggesting that overall Torso levels are higher in the absence of functional ESCRT-0 (**Figure S2C**). Together these perturbations validate that the dynamic range of the Torso::pYtag system is appropriate for sensing both increases and decreases in Torso activity from the wild-type level.

### Dynamics of Torso activity differ from downstream ERK activity

Having shown that Torso::pYtag is a reliable biosensor for endogenous Torso activity, we next sought to use Torso::pYtag to investigate how the ERK gradient is established. Focusing on the approximately 1-hour window of nuclear cycles (NC) 11 to early 14 when Torso activity drives terminal ERK signaling at the poles, we compared Torso::pYtag activity to two established techniques for measuring this gradient at downstream signaling nodes: immunostaining for doubly-phosphorylated, active ERK (dpERK) downstream of Ras (Gabay et al., 1997), and the miniCic biosensor that reflects phosphorylation of the ERK substrate and transcriptional repressor Capicua (Moreno et al., 2019) (**Figure 3A**). Previous characterization of ERK activity in the embryo using the dpERK antibody showed that an initially broad and shallow gradient of ERK spanning 40% of the embryo’s length sharpens, increases in maximum amplitude, and retracts towards the poles over time (Coppey et al., 2008). We replicated these findings by staining and quantifying dpERK in wild-type embryos across the anterior-posterior axis (**Figure 3B, 3C** - center). Similarly, the live miniCic biosensor shows that the domain of ERK kinase activity spans a similar length and retracts toward the poles over time (**Figure 3B, 3C** – right). We note that miniCic has a high sensitivity to ERK activity but a low dynamic range, and so it is a good indicator of the size of the domain of ERK kinase activity but cannot report on the amplitude at the poles where ERK activity is highest.

**Figure 3:**
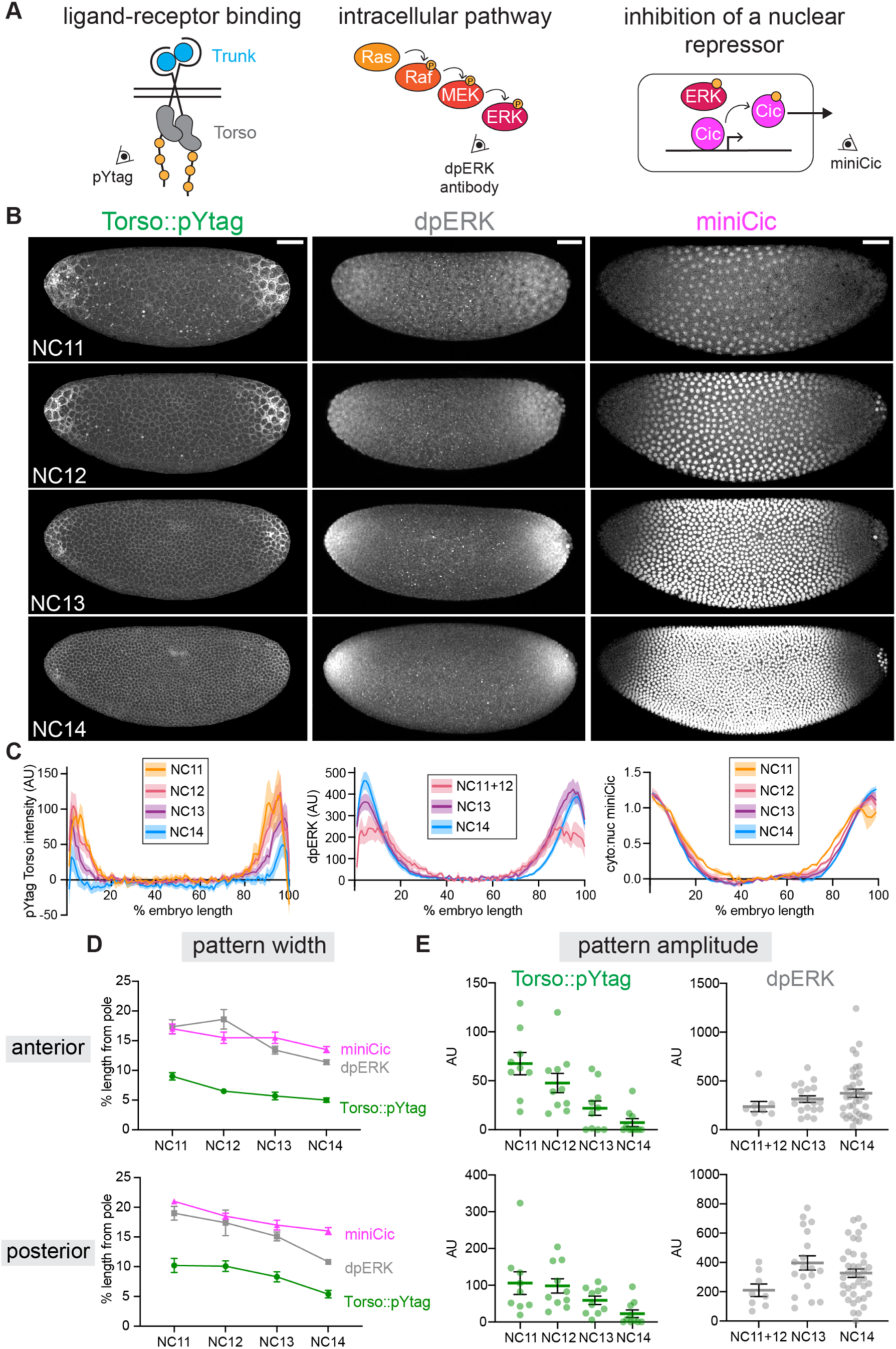
Dynamics of Torso activity differ from downstream ERK activity. (A) Imaging of terminal signaling at different nodes of the pathway: receptor activation (Torso::pYtag), intracellular pathway activity (dpERK antibody), and inhibition of a nuclear repressor by ERK kinase activity (miniCic). (B) Images show embryos in NC11-14 visualized at each node. Left: Torso::pYtag images taken from a single embryo over time. Center: WT embryos stained for dpERK. Right: miniCic::mCherry images taken from a single embryo over time. Scale bar: 50 μm. (C) Quantification of each node across the anterior-posterior axis in each nuclear cycle. Torso::pYtag (left) and dpERK (center) intensity values are normalized to the center of the embryo. miniCic (right) cytoplasmic:nuclear ratio is normalized to the highest and lowest values. Line shows mean ± s.e.m. of n = 10 Torso::pYtag embryos, n = 3 miniCic embryos, n = 8 (NC11+12), 19 (NC13), 39 (NC14). (D) Pattern width (measured as % embryo length from each pole) and (E) pattern amplitude over time for each embryo quantified in (C). Mean ± s.e.m.

Based on these measurements of ERK activity, we predicted that Torso::pYtag would be active in the same region as ERK and show similar features: decreasing size of the signaling domain with constant or increasing signaling amplitude. However, when we live imaged Torso::pYtag over NC11-14 and quantified Torso::pYtag intensity across the anterior-posterior axis, we observed an unexpected pattern of Torso activity (**Figure 3B, 3C** – left, **Movie S1**): a pronounced decrease in the intensity of Torso::pYtag over time across the entire pattern, and a spatial domain of Torso activity that was severely restricted to the poles, particularly in NC13-14. These data suggest that during terminal patterning, a broad and sustained ERK activity gradient is produced by a narrow and decreasing receptor-level stimulus.

To better compare dynamics across time and measurement type, we used two metrics of activity. First, we quantified the “pattern width” by determining the position (measured as % embryo length from the pole) at which each readout reached its half maximum intensity (**Figure 3D**). The pattern widths of Torso::pYtag, dpERK, and miniCic all decreased by about 5% of the embryo’s length between NCs 11 and 14, indicating that both receptor-level and downstream pathway activity occur over successively smaller domains, likely driven by ligand trapping at the poles (Casanova and Struhl, 1993; Coppey et al., 2008). However, the dpERK and miniCic patterns were always wider than the Torso::pYtag pattern by roughly 10% (50 μm) of the embryo length. The observation of a narrow pattern of Torso activity giving rise to a broader ERK pattern would be consistent with diffusion of active ERK pathway components in the syncytial embryo (Ho et al., 2023).

We then assessed “pattern amplitude” for both Torso::pYtag and dpERK by averaging the signal intensity at 5-10% (anterior pole) and 90-95% (posterior pole) egg length of the embryo. At both the anterior and posterior poles, Torso::pYtag amplitude decreased to ∼10% of its NC11 intensity by NC14, whereas dpERK amplitude doubled over this same time period (**Figure 3E**). This observation strongly suggests that substantial decreases in Torso signaling amplitude are not accompanied by decreases in downstream ERK activity, a surprising outcome given that downstream pathway activity is known to depend on continued upstream input (Keenan et al., 2020).

We confirmed that this decrease in Torso::pYtag amplitude over time was not due to bleaching by comparing the pattern amplitude of embryos imaged over all nuclear cycles with embryos imaged only in NC14. There was no difference in the pattern amplitude between these conditions (**Figure S3A**). We also observed a similar decrease in pYtag amplitude in two other versions of the pYtag: embryos lacking the 3xP3-DsRed (**Figure S3B**) and embryos with a red version of the biosensor (**Figure S3C**). We did not observe a decrease in total Torso protein using an endogenously-tagged Torso (Torso::sfGFP) (**Figure S3D**), consistent with previous measurements that show consistently high receptor levels throughout this period (Casanova and Struhl, 1989; Sprenger et al., 1993). Therefore, we conclude that Torso activity decreases over time in contrast to increasing ERK activity over the same window.

### Negative feedback attenuates Torso activity

Despite some changes in steepness and amplitude, the ERK gradient is remarkably stable over a developmental period in which major changes are occurring in number of nuclei, zygotic gene expression, and, now we find, Torso activity. These observations suggest that the ERK gradient might be stabilized by feedback. Feedback regulation is a common mechanism for shaping gradients amidst developmental noise (Shilo and Barkai, 2017). Feedback is also mechanistically plausible, as phosphatases can both serve as targets of ERK phosphorylation and as regulators of receptor activity (Neben et al., 2019; Wang et al., 2002; Wang et al., 2007). However, feedback control during terminal patterning has not been fully explored. The pYtag offers a novel opportunity to test for such feedback directly *in vivo*.

To test whether the decrease in Torso activity is due to negative feedback from ERK, we blocked ERK activation downstream of Torso using an RNAi targeting *kinase suppressor of ras* (*ksr*) (**Figure 4A**). Ksr is a scaffold protein necessary for signal propagation through the Ras/ERK pathway (Roy et al., 2002). In the absence of Ksr, Torso can be activated by ligand, but there is no activation of ERK and thus no opportunity for induction of feedback. In embryos expressing maternal *ksr* RNAi, miniCic was nuclear throughout the embryo indicating strong suppression of ERK activity (**Figure S4A, S4B**). While RNAi control embryos expressing only the maternal Gal4 showed decreasing Torso::pYtag activity over time, *ksr* RNAi embryos showed remarkably strong Clover-ZtSH2 membrane recruitment at the poles in all nuclear cycles (**Figure 4B**, **Movie S2**). The difference was most striking in NC14, as shown by comparison of Torso::pYtag intensity along the anterior-posterior axis in NC14 (**Figure 4C**). *ksr* RNAi embryos showed increasing pattern amplitude over NC11-14 with significantly higher pattern amplitude than no-RNAi controls in NC14 (**Figure 4D**, **S4C**). Therefore, in the absence ERK activity, Torso phosphorylation remains high, strongly suggesting a role for ERK pathway-dependent negative feedback in attenuating Torso activity. Notably, the pattern width of *ksr* RNAi embryos however was unchanged from controls (**Figure 4E**, **S4D**), suggesting that the spatial domain of Torso activity is regulated independently from the amplitude of Torso activity by ligand distribution.

**Figure 4:**
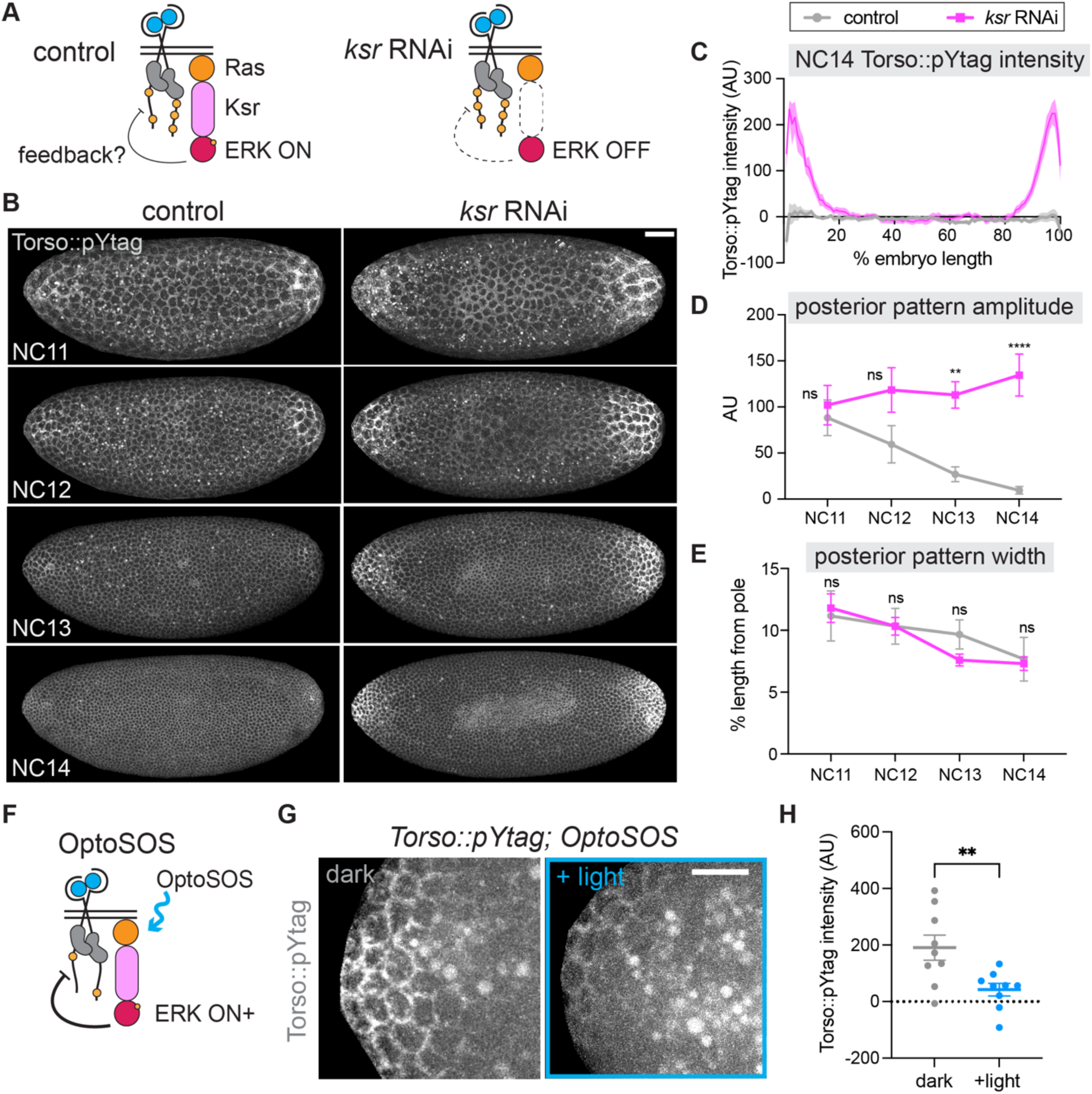
Disrupting ERK pathway activity impacts Torso activity, revealing negative feedback. (A) Diagram shows that removal of the scaffold protein Ksr inhibits ERK activity, preventing negative feedback on Torso without impacting Torso activation by Trunk ligand. (B) Images taken from a single control and *ksr* RNAi embryo expressing Torso::pYtag over NC11-14. Scale bar: 50 μm. (C) Quantification of the anterior-posterior intensity of Torso::pYtag in control and *ksr* RNAi embryos in NC14. Line shows mean ± s.e.m. of n = 11 (control), 10 (*ksr*) embryos. (D) Posterior pattern amplitude and (E) posterior pattern width over time for each embryo quantified in (C). Mean ± s.e.m. Significance by 2-way ANOVA with multiple comparisons test. (F) Diagram shows how OptoSOS induction of Ras activity increases ERK activity and induces negative feedback on Torso. (G) Images show Torso::pYtag at the anterior pole of NC13 in unilluminated (dark) and illuminated (+light) OptoSOS embryos. The bright circles in the images are autofluorescent yolk. Scale bar: 20 μm. (H) Quantification of Torso::pYtag intensity at the anterior pole in each condition. Mean ± s.e.m. n = 9. Unpaired t test.

We then reasoned that if there is ERK pathway-dependent feedback on Torso activity, we should be able to further decrease Torso signaling by increasing ERK pathway activity. To do so, we turned to an optogenetic technique to activate ERK and measure any resulting changes in Torso::pYtag activity (**Figure 4F**). OptoSOS is an optogenetic tool for activating the Ras/ERK pathway that functions by recruiting the catalytic domain of SOS to the membrane using the iLID-SspB dimerization system (Johnson et al., 2017; Toettcher et al., 2013). We generated embryos expressing both Torso::pYtag and OptoSOS and stimulated the anterior poles of embryos with blue light from NC10 to early NC13 (approximately 25-30 minutes). Consistent with our prediction, OptoSOS embryos stimulated with blue light had significantly lower Torso::pYtag activity than unstimulated OptoSOS embryos in NC13 (**Figure 4G,H**). The posterior poles of these same embryos, which were unilluminated in all cases, showed no difference (**Figure S4E**). There was also no decrease in Torso::pYtag intensity in embryos lacking OptoSOS but illuminated with the same blue light stimulus, again showing that bleaching is not the cause of this change (**Figure S4F**). Therefore, there is potent negative feedback on Torso phosphorylation state from the ERK pathway that responds to both decreases and increases to downstream ERK pathway activity.

### A local domain of Torso activity produces a longer range ERK gradient

We then considered a second surprise from the comparison of Torso and ERK activity: the spatial pattern of Torso activity is quite different from that of the ERK gradient it produces. It has long been an open question whether the ERK gradient derives from graded Torso activity or whether it arises by diffusion of active signaling components away from a local domain of Torso activity (Casanova and Struhl, 1989). We previously showed that diffusion in the syncytial embryo is sufficient to blur a sharp boundary of photoconverted fluorescent proteins into a gradient extending 50 μm within minutes, and also to create a broad terminal signaling gradient downstream of a narrow, all-or-none optogenetic Ras input (Ho et al., 2023). However, determining whether this diffusion is the primary contributor to the endogenous ERK gradient requires directly assessing the pattern of receptor activity, which is now possible with pYtags.

Our measurements of Torso::pYtag, dpERK, and miniCic show that the domain of ERK activity extends beyond the major domain of Torso activity about 50 μm or 10% of the embryo length in all nuclear cycles (**Figure 3D**), consistent with a key role for diffusion. To examine this discrepancy more closely, we asked if ERK activity outside the domain of Torso activity is functionally sufficient to induce target gene expression. Tailless (*tll)* is an essential ERK target gene expressed at low levels of ERK activity (Pignoni et al., 1990). *Tailless* transcription is a useful metric of active ERK signaling because it responds rapidly and reversibly (within 5 minutes) to optogenetic ERK pathway activation (Ho et al., 2023; Keenan et al., 2020). We generated embryos expressing Torso::pYtag together with a fluorescent MCP protein for visualizing transcription of MS2-tagged transcripts and MS2 stem loops in the endogenous *tll* 5′ untranslated region (*tll*-MS2) (Garcia et al., 2013; Keenan et al., 2022). Imaging of Torso::pYtag and *tll*-MS2 revealed that *tll* transcriptional bursts occur outside the domain of active Torso::pYtag (**Figure 5A**). *tll* bursts were present up to 50 μm outside the major domain of Torso::pYtag activity (**Figure 5B**). Although we cannot exclude that there is low level Torso activation undetected by the biosensor driving this *tll* expression, our data are more consistent with the model of ERK gradient formation where a local domain of Torso activity at the poles is blurred to produce a longer-range gradient of ERK activity (**Figure 5C**).

**Figure 5:**
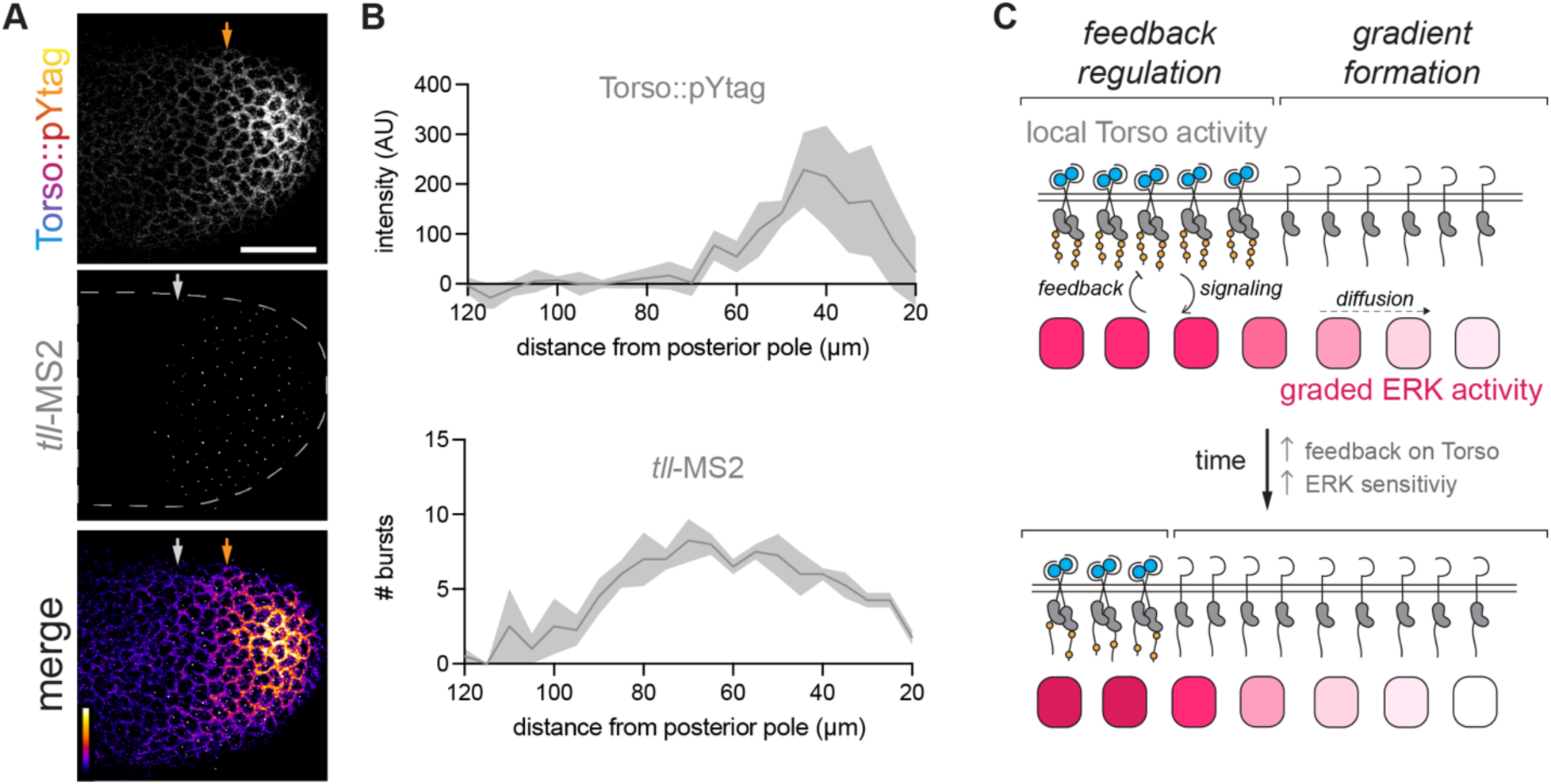
A local domain of Torso activity produces an extended domain of ERK signaling. (A) Images of a NC13 embryo showing Torso::pYtag together with *tll* MS2 transcriptional bursts at the posterior pole. Arrows indicate the boundaries at each pole of the Torso ON (orange) and *tll* expressing (white) regions. Scale bar: 50 μm. (B) Quantification of Torso::pYtag intensity and number of *tll-*MS2 bursts measured from the posterior pole. Line is mean ± s.e.m. n = 4 embryos. (C) Model of terminal patterning. A local domain of Torso activity creates a gradient of ERK activity by diffusion. In the region of Torso activity, ERK pathway activity feeds back on Torso activity to attenuate receptor activity.

## Discussion

Here, we have applied and validated pYtags, a new class of RTK biosensors, for detection of endogenous RTK activity in developing tissues. We engineered pYtags for three *Drosophila* RTKs and confirmed that these biosensors display expected patterns of receptor activity in diverse developmental contexts: Torso in the syncytial embryo, EGFR in the epithelial follicle cells of the egg chamber, and Btl in migrating tracheal tip cells. Using Torso as a model RTK, we characterized the dynamics of Torso activity that determine the terminal ERK gradients for the first time. Through comparison of Torso activity with downstream ERK, we identified a role for negative feedback in attenuating receptor activation and investigated the spatial discrepancy between the Torso and ERK activity domains.

### Endogenous biosensors of RTK activity

Direct detection of RTK activity has the potential to reveal many previously obscured features of receptor signaling, and our work shows that pYtags are well-suited for these efforts. First, pYtags reveal subcellular activation of RTKs, as shown by terminal Torso activity within the syncytial embryo and apical activation of EGFR in the follicle cells. We similarly expect pYtags will be useful for assessing subcellular RTK activation within migrating cells undergoing chemotaxis (Gong et al., 2024; Haugh et al., 2000; Janssens et al., 2010; Jha et al., 2025). Second, pYtags allow quantitative measurements of receptor-ligand interactions which can be used to directly visualize morphogen gradients and then compare receptor activity with downstream signaling. Using this approach in the context of the canonical terminal pattern, we unexpectedly found that the dynamics of Torso and ERK activity are not correlated in the early embryo in space or time, highlighting that receptor activity cannot be directly inferred from downstream signaling activity. Finally, we envision that pYtags will be useful for distinguishing the roles of specific RTKs in tissues where inputs to multiple RTKs contribute to the same processes (Duchek and Rørth, 2001; Duchek et al., 2001; Freeman, 1996).

pYtags have broad applicability because in principle they can be applied to any RTK of interest without receptor-specific design changes. We have thus far validated three *Drosophila* pYtags, but pYtags can be readily applied to any of the 20 *Drosophila* RTKs and, we expect, RTKs in other organisms (Sopko and Perrimon, 2013). Our work highlights the usefulness of pYtags but also reveals a few challenges for their use *in vivo*. First, low expression of RTKs may be a challenge for detection of the biosensor in some tissues. Some RTKs are known to be expressed at low levels (Janssens et al., 2010; Sapir et al., 1998), and ZtSH2 membrane recruitment might be particularly challenging to detect in these contexts. Second, we did not find any evidence that the homology of the ITAM and Zap70 components to endogenous proteins caused any non-specific interactions of the pYtag, but we can’t exclude that there may be challenges in other tissues (Torres et al., 2023). Finally, while the interaction between the ITAM and ZtSH2 is highly specific (Isakov et al., 1995), there is limited specificity for tyrosine phosphorylation by tyrosine kinases (Hunter, 2009). In principle, another tyrosine kinase could phosphorylate the ITAM and drive membrane recruitment of ZtSH2. Our data suggest that the Torso::pYtag may detect the global increase in activity of another tyrosine kinase during NC13 and NC14 in the *Drosophila* embryo, since an ITAM-less control receptor does not display a similar global increase in ZtSH2 membrane recruitment (Loncar and Singer, 1995). Alternatively, it is possible that the baseline pYtag translocation reflects a low level of ligand-independent RTK kinase activity. It is critical that any application of pYtags includes scenarios where ligand/receptor activity is zero (such as comparisons to *tsl* mutants for Torso activity or comparisons to the ventral side of the egg chamber for EGFR activity) to establish the baseline level of pYtag translocation.

### An updated model for terminal patterning

Our characterization of Torso::pYtag revealed two surprises about terminal patterning made possible by the ability to directly compare receptor activity with downstream ERK pathway activity *in vivo*. The first surprise was that the spatial domains of Torso and ERK activity are distinct: the Torso activity gradient is sharper and more restricted to the poles, whereas ERK activity is strongly graded and extends farther toward medial positions. We measured approximately a 50 μm gap between the boundaries of Torso activity and *tailless* transcription. This measurement was remarkably consistent with our previous findings that ERK activity extends approximately 50 μm beyond a boundary of optogenetic Ras activation and that diffusion in the syncytial embryo is sufficient to blur a sharp boundary along this length scale within minutes (Ho et al., 2023). Thus, both natural (by Torso) and synthetic (by OptoSOS) activation of local Ras result in longer range formation of an ERK gradient likely due to diffusion of active signaling components. These findings provide evidence supporting a model of terminal patterning where gradient formation occurs primarily downstream of local receptor activity.

The second surprise was a striking difference in the trajectories of Torso and ERK amplitude over time. While it was previously known that ERK activity maintains a high or increasing amplitude over NCs 11 through 14 (Coppey et al., 2008), we found that Torso activity decreases over the same hour-long interval. This result was surprising because downstream pathway activity is known to respond rapidly to changes in receptor-level inputs. Previous work showed rapid (within 5 minutes) cessation of *tailless* target gene transcription upon removal of optogenetic activation of Ras (Keenan et al., 2020). We hypothesize that ERK activity remains high as Torso activity falls because ERK becomes more sensitive to Torso inputs in later nuclear cycles. This change in sensitivity might be driven by the onset of zygotic expression of Torso-to-ERK pathway components, physical changes to the embryo, or competition for other factors. As an example of how other factors can modulate ERK sensitivity, it is known that anterior Bicoid and dual specificity phosphatases compete for binding to ERK, resulting in higher levels of active ERK in the anterior than the posterior (Kim et al., 2010).

Our experiments reveal ERK pathway-dependent feedback on Torso phosphorylation state: by engineering embryos with either increased or decreased ERK signaling, we observe changes in Torso-pYtag activity within as little as ∼30 min. RTKs are known to be regulated by a variety of phosphatases, but evidence for ERK-dependent regulation of these phosphatases is limited (Neben et al., 2019; Prahallad et al., 2012). Alternatively, there is evidence for direct phosphorylation of RTKs by ERK which might influence receptor phosphorylation by modulating its kinase activity, recruiting phosphatases, or affecting receptor ubiquitination and trafficking (Li et al., 2008; Takishima et al., 1991; Zakrzewska et al., 2013). We must also emphasize that the discovery of ERK-to-Torso feedback does not prove a complete feedback loop, where the new, altered Torso activity state further changes ERK activity. It remains possible that the phosphorylation state of the ITAM does not reflect the phosphorylation state of all tyrosines on the receptor, and ERK-to-Torso feedback might serve to tune receptor activity in a more nuanced manner. Future biochemical analyses will address these additional mechanistic questions.

Nevertheless, it is tempting to speculate about the roles that feedback regulation might play in an updated model of terminal patterning in which a highly regulated signaling system maintains a simple gradient (**Figure 5D**). Torso receptors are activated in a narrow domain near the poles by Torso ligand. Over time, the Torso-active domain retracts towards the poles, likely due to ligand trapping, and its amplitude decreases in response to ERK pathway-dependent negative feedback. This feedback likely plays a critical role in maintaining appropriate levels of ERK signaling for downstream gene expression because corresponding increases in ERK pathway sensitivity might otherwise inappropriately expand these patterns. Diffusion of intracellular signaling components then transforms this narrow receptor pattern into a longer-range gradient of ERK. Thus, the amplitude of the ERK gradient and its spatial range are independently controlled by two processes, receptor-level feedback and intracellular diffusion, respectively. Future work will address key outstanding questions about the mediators of feedback, the mechanisms changing ERK pathway sensitivity, and the diffusive components of the Ras/ERK pathway. However, the insights here are made possible by the ability to directly detect receptor activity using pYtags, and we look forward to similar insights in other patterning systems.

## Materials and Methods

### Fly stocks

Flies were cultured using standard methods at 25°C. All pYtag lines and Torso::sfGFP were made in this paper using CRISPR/Cas9 editing described below. Other stocks used include *tub-miniCic::mCherry* (Moreno et al., 2019), *tub-miniCic::NeonGreen* (gift of Romain Levayer), *MCP::mCherry* (gift of Mike Levine), *67;15* (*P{matα-GAL-VP16}67; P{matα-GAL-VP16}15*, Bloomington Drosophila Stock Center (BDSC), #80361), y^1^w^118^ (BDSC #6598), *trk^1^* (BDSC #1212), *tsl^4^* (BDSC #3289), *Hrs* RNAi (BDSC #34086), *ksr* RNAi (BDSC #41598), *UAS-OptoSOS* (Johnson et al., 2017), *tll-MS2* (Keenan et al., 2022)*, CAAX-mCherry*, *His2Av-GFP* (gift of Eric Wieschaus), *btl-Gal4*, and *UAS-CD4-mIFP* (BDSC #64183).

Recombination of *Torso::pYtag* and *EGFR::pYtag* on the second chromosome with *miniCic, P{matα-GAL-VP16}67, or MCP::mCherry* was confirmed using visible markers (mini-white and DsRed positive) and functional validation. Recombination of *Torso::pYtag* with *trk^1^* was confirmed using visible DsRed together with maternal lethality of the *trk^1^* allele. To produce viable *tsl^4^* homozygous females, a chromosome with *tsl^4^* recombined with *P{matα-GAL-VP16}15* was used in trans with *tsl^4^* (Johnson et al., 2020).

For optogenetic experiments, the *67;15* Gal4 driver was used for maternal expression of OptoSOS. *Torso::pYtag, 67;15* females were crossed with *Torso::pYtag, UAS-OptoSOS* males. The resulting *Torso::pYtag, 67/Torso::pYtag; 15/UAS-OptoSOS* females were used for the experiment and placed in the dark at room temperature for at least two days prior to the experiment. For RNAi experiments, the *67;15* Gal4 driver was used for maternal expression of RNAis. *Torso::pYtag,67;15* females were crossed with *Torso::pYtag, miniCic::mCherry; RNAi* males containing the desired RNAi or no RNAi for controls. The resulting *Torso::pYtag, 67/ Torso::pYtag, miniCic::mCherry; 15/ RNAi* females were used for the experiment and placed at 18°C. Successful knockdown was confirmed by assessing the change in ERK signaling using miniCic::mCherry.

### Plasmid generation

All cloning was performed using InFusion cloning. Linear DNA fragments were amplified by PCR using HiFi polymerase (Takara Biosciences), digested with DpnI to remove template DNA (NEB), and isolated by gel electrophoresis and purification (NucleoSpin columns, Takara Biosciences). Fragments were assembled using InFusion assembly and transformed into Stellar competent *E. coli* cells (Takara Biosciences). Plasmids were purified using a Qiagen miniprep kit and validated using whole plasmid sequencing (Plasmidsaurus).

Guide sequences for C-terminal tagging of RTKs were designed using the FlyCRISPR Optimal Target Finder or taken from published work (Gratz et al., 2014; Revaitis et al., 2020). The guide sequences (with PAM sequence underlined) were: Torso AGCAATGACTATTAATTCAA<u>AGG</u>, EGFR GAAACCGCAACACGGAGACG<u>AGG</u>, and Btl GGTGTACTGATATCTAAGCG<u>GGG</u>.

Oligos containing the guides (**Table S1**) were annealed and ligated using T4 ligase into BbsI digested pU6-BbsI-chiRNA vector (Addgene #45946) (Gratz et al., 2013).

To produce the homologous repair construct for pYtag insertion, a 3xCD3ε-P2A-Clover- ZtSH2 construct was assembled from the 3xCD3ε ITAM sequence of pHR EGFR-CD3ε- FusionRed (Addgene #188626) and the ZtSH2 sequence from pHR iRFP-ZtSH2 (Addgene #188627) (Farahani et al., 2023) and then inserted into pScarlessHD-DsRed (Addgene #64703) to produce pScarlessHR 3xCD3ε-P2A-Clover-ZtSH2-DsRed. A red variant of this pYtag repair construct was made by swapping Clover with mScarlet-I, and an ITAM-less version was made by removing the 3xCD3ε. 5’ and 3’ homology arms (1 kb) flanking the stop codons of *torso*, *Egfr*, and *btl* were amplified from genomic DNA that had been extracted from *nos-Cas9* flies (BDSC #78781) using primers in **Table S1**. The homology arms were then inserted into the 5’ and 3’ ends of the pYtag repair construct. To prevent retargeting of the guide to the edited locus, synonymous mutations were introduced to the repair sequence either with site directed mutagenesis (EGFR) or in the primers used to amplify the homology arms (Btl). The Torso guide sequence was not altered because it would be destroyed by successful insertion.

To produce the homologous repair construct for C-terminal tagging of Torso with superfolder GFP (sfGFP), the pYtag portion of pScarlessHR Torso-3xCD3ε-P2A-Clover-ZtSH2-DsRed was replaced with sfGFP to produce pScarlessHR Torso-sfGFP-DsRed. A Gly-Gly-Gly linker was included between Torso and sfGFP.

### *Drosophila* CRISPR/Cas9 editing

The homology arms and gRNA plasmids were co-injected into *nos-Cas9* flies by BestGene Inc and screened for 3xP3-DsRed insertion. Candidate lines were genotyped by PCR with primers flanking the homology arms (**Table S1**) and then the linear amplicons were fully sequenced to confirm successful editing (Plasmidsaurus). The pYtag is ∼1.8 kb and the total insertion is ∼3.5 kb. Most lines used retained the 3xP3-DsRed, but when noted the DsRed was removed by crossing to *nos-PBac* flies (Keenan et al., 2022). The resulting –DsRed lines were validated by PCR and sequencing.

### Sample preparation for live imaging

Female flies of the desired maternal genotype (**Table S2**) were placed in a cage with standard apple juice plates and yeast paste. Mothers were always homozygous for the desired pYtag allele. For MS2 experiments, virgin females were mated with homozygous MS2 males. For all other experiments, non-virgin females were mated either with their siblings or wild-type (*y^1^w^118^*) flies. The cage was placed at room temperature for at least 2 days before collection for all experiments except for RNAi experiments in which the cage was placed at 18°C to increase expression from the *67;15* Gal4 driver. For OptoSOS experiments, the cage was placed in the dark.

For Torso::pYtag imaging, embryos collected 0-3 h post egg lay were manually dechorionated with double-stick tape and then mounted on their lateral side between a semi-permeable membrane and a coverslip in a 3:1 mix of halocarbon 700/27 oil. For Btl::pYtag imaging, embryos at roughly stage 13 were dechorionated in bleach and placed in a glass bottom dish with water. For EGFR::pYtag imaging, ovaries were dissected from mated females placed on wet yeast for at least 3 days, individual ovarioles were separated, mounted in a thrombin (GE Healthcare) and fibrinogen (Sigma Aldrich) clot, and live imaged in Schneider’s insect media supplemented with FBS and insulin (Sigma Aldrich) (Wilcockson and Ashe, 2021).

### Live microscopy

Torso::pYtag imaging was done on a Nikon A1R scanning confocal microscope equipped with a 25 mm galvano scanner. At 40x, the entire embryo can fit within the field of view. For imaging at 60x, only the posterior was imaged. Images were acquired once each nuclear cycle (NC10 - early NC14) with 4x averaging. Roughly a 30 μm z depth (1 μm step size) was taken, spanning from the surface of the embryo to a z plane in which the pole cells were visible to ensure the entire length of the embryo was captured. If the pole cells could not be captured without significantly increasing the z depth, a final image was taken that included the pole cells to determine the length of the embryo. Any embryos visibly injured by dechorionation or exhibiting uneven cell cycles were excluded. To capture higher time-resolution dynamics with 3-minute intervals at 40x, acquisition speed was increased by cropping the imaging area and reducing to 2x averaging. Torso::sfGFP imaging was done similarly at 40x. EGFR::pYtag imaging was done similarly at 40x with 4x averaging and 1 μm z steps to the midplane of the stage 10 egg chamber. Images of *miniCic::mCherry ; His2Av-GFP* embryos were taken similarly at 40x with 2x averaging and 2 μm z steps. Btl::pYtag imaging was done on a Nikon Ti2 microscope equipped with a Yokogawa spinning disc (CSU-W1) and Hamamatsu BT Fusion sCMOS detector at 40x. Z-stacks were denoised using Nikon Elements software.

### Optogenetic stimulation

Optogenetic stimulation was performed at room temperature at 40x on a Nikon Eclipse Ti microscope with a Yokogawa CSU-X1 spinning disk and an iXon DU897 EMCCD camera equipped with a Mightex Polygon digital micromirror device and an X-Cite LED 447-nm blue LED. The anterior pole of an embryo was stimulated with a 0.6 sec pulse of 23 mW/cm^2^ blue light applied every 30 sec. Under these conditions, the SSPB-tagRFP-SOScat component of OptoSOS strongly relocalized to the membrane. From previous optogenetic work, it is known that this strong membrane localization of OptoSOS represents a high level of ERK activity (sufficient to induce high ERK target genes such as *huckebein*) (Ho et al., 2023). Stimulation was performed from NC10-13 (25-30 min), and then a full z-stack image of Torso::pYtag was captured at the final timepoint in NC13.

### Embryo fixation and staining

Wild-type (*yw*) or Torso::pYtag embryos were bleached, fixed in an 8% paraformaldehyde/heptane interface, and devitellinized with methanol as described previously (Coppey et al., 2008). For dpERK immunostaining, embryos were rehydrated in PBS + 0.02% Triton-X (PBST), treated with Image-iT signal enhancer (Invitrogen), and then blocked with 5% Normal Goat Serum (NGS), 10% BSA in PBST. Embryos were treated with primary Rabbit anti-dpERK antibody (Cell Signaling #4370) at 1:100 in PBST with 5% NGS, 1% BSA overnight at 4°C. After washing, embryos were treated with secondary Goat anti Rabbit 647 (Invitrogen) at 1:500 and DAPI for nuclear staining in PBST with 10% NGS, 1% BSA for 2 hours at room temperature. Following final washes, embryos were mounted using Aqua-Poly/Mount (Polysciences). HCR v3.0 RNA *in situ* staining was performed according to (Choi et al., 2018) using initiators and buffers from Molecular Probes. Probes for *tll* (NM_079857.4) and *hkb* (NM_079497.4) were either ordered from Molecular Probes or designed using insitu_probe_generator software (Kuehn et al., 2022) and synthesized as a oligo pool by IDT. DAPI was added in one of the final wash steps to stain nuclei. Fixed and stained embryos at the desired stage were imaged at 20x on a Nikon A1R scanning confocal microscope by taking a z stack from the surface to the medial plane.

### Torso::pYtag quantification

Images were processed using ImageJ. The green autofluoresence of the vitelline membrane was manually segmented out of all Torso::pYtag images. Then, a maximum z-projection was produced for each timepoint. Because there was not clear cytoplasmic localization of ZtSH2 in regions without active Torso, Clover-ZtSH2 intensity was used as the measure of Torso activity. To quantify the anterior-posterior profile of Torso activity, a line with a width of 40 pixels (∼35 μm) was drawn from the anterior pole to the posterior germ cells and Clover-ZtSH2 intensity was measured along the line. For a given embryo, the same line was used in all nuclear cycles. To compare this line scan between embryos, ZtSH2 intensity was normalized by subtracting the ZtSH2 intensity from the center, Torso-OFF, region of the embryo. In some embryos that were not mounted level to the coverslip, the background was uneven between the anterior and posterior sides of the embryo. In these cases, a background intensity was determined for the anterior and posterior halves of the embryo separately. Next, the line scan was binned into 100 bins, each representing 1% of the embryo length, and the average intensity in each bin was determined. In cases where the total embryo length was longer than the captured length as described above, the line was drawn based on the total embryo length and intensity values representing the unimaged length (usually 1-2% embryo length at each end) were excluded. Torso::sfGFP was quantified similarly but was not normalized to the center of the embryo. For imaging at 60x of the posterior pole, the line scan used was 80 pixels (∼40 μm) and binning was performed at 5 μm intervals.

To determine the “pattern width” of Torso::pYtag in each embryo and at each nuclear cycle, the absolute maximum value at each pole was determined from the binned and averaged line scan. Then the position (measured in % embryo length from the pole) at which the Torso::pYtag value dropped permanently below 50% of the maximum value was set as the pattern width. In cases where there was no detectable peak of Torso::pYtag at the pole, the pattern width was not calculated. To determine the “pattern amplitude” of Torso::pYtag, the intensities (again from the binned and averaged line scan) were averaged over 5-10% and 90-95% embryo length. This position was chosen because it was difficult to capture the last 5% of the embryo length due to the curve of the embryo at the poles. If the calculated pattern amplitude was a negative number, it was set to 0.

To quantify the background Torso::pYtag intensity in the center of the embryo where Torso activity is off, a 50 x 50 μm box was drawn in the center of the maximum-projected image and positioned to avoid bright yolk granules. The mean intensity within this box was measured. To normalize background fluorescence across nuclear cycles, the intensity was divided by the NC11 intensity.

Quantification of Torso::pYtag in optogenetic experiments required a different approach because the spinning disc microscope used produced a noisier image with a much smaller field of view. First, rolling ball background subtraction with a radius of 50 was applied to each z slice and then a maximum intensity projection was produced. Then, a 25 x 25 μm box was drawn at the anterior pole of the embryo in the region the Torso::pYtag was on, and a background box of the same size was drawn farther from the pole along the edge of the embryo where Torso::pYtag was off. Autofluorescent yolk granules were avoided. Then the mean intensity of the background box was subtracted from the mean intensity of the Torso::pYtag box.

### EGFR::pYtag quantification

A maximum z projection was made with the 5 z-slices at the medial plane of the embryo where the oocyte nucleus and follicle cell cross sections were visible. Apical membrane enrichment of the EGFR::pYtag was quantified by drawing a line with a width of 5 pixels (∼3 μm) along the apical surface of the follicle cells and then a parallel line was drawn in the apical cytoplasm (between the membrane and nucleus). The Clover-ZtSH2 intensity was measured along both lines, the membrane intensity was divided by the cytoplasmic intensity at each pixel, and the result was binned over 3 pixels.

### Btl::pYtag quantification

To quantify activity of the Btl::pYtag, z-stacks were maximum projected. Then the membrane localization of the Btl::pYtag was determined by assessing colocalization of Btl::pYtag and the membrane marker CD4::mIFP. The stalk-to-tip axis of each tracheal branch was tiled with 7-9 boxes measuring 4.5 x 4.5 μm. Within each box, the Pearson’s correlation was calculated between the Btl::pYtag and membrane pixel intensities.

### miniCic quantification

For a simple assessment of ERK activity using miniCic, the proportion of the embryo in an ERK-off state was determined by measuring the proportion of total embryo length with nuclear miniCic. If there is no terminal ERK activity, as in a *tsl* mutant, this metric equals 1. For a quantitative profile of miniCic localization across the embryo, nuclei from images containing Histone-GFP and miniCic::mCherry were segmented using Cellpose (Stringer et al., 2021), and a dilation and subtraction were applied to the segmented nuclei to isolate the immediately surrounding cytoplasmic region. Then the cytoplasmic:nuclear miniCic intensity ratio was measured along the anterior-posterior axis, binned into 50 equal-sized bins, and normalized so that each embryo spanned from 0 (in the center of the embryo) to 1 (at the anterior pole). miniCic pattern width was calculated similarly to Torso::pYtag.

### dpERK quantification

To measure dpERK intensity across the anterior-posterior axis, images of fixed and anti-dpERK stained embryos were first sorted by nuclear cycle based on the nuclear density. Embryos in late NC14 or undergoing mitosis were excluded. A maximum z projection was made and then a 20 pixel (∼30 μm) wide line was drawn from the anterior tip to posterior pole. The intensity of dpERK along this line was measured. Similar to quantification of Torso::pYtag, the line scan values were normalized by subtracting the intensity at the center (ERK OFF) region of the embryo, binned into 100 bins each representing 1% of the embryo length, and then averaged in each bin. dpERK pattern width and pattern amplitude were calculated similarly to Torso::pYtag.

### *tll*-MS2 quantification

Maximum projections of *tll-*MS2 bursts were thresholded and analyzed to identify particles. The position of each particle was determined relative to the posterior pole and burst counts were binned in 5 μm bins.

### *tll* and *hkb* quantification

To quantify HCR staining of *tll* and *hkb* mRNA, a line with width 20 pixels (15 μm) was drawn along the anterior-posterior axis of the maximum intensity z projection. Then the intensity values were binned into 100 equal bins and normalized over a 0 to 1 range for each embryo.

### Statistical analysis

All statistical analysis was performed in Prism 10 (GraphPad). Details for statistical tests used can be found in the figure legends. Figure legends indicate the number of embryos for each condition (n). In dot plots, each dot is one embryo. All graphs show the mean ± s.e.m. Significance was defined as *P<0.05, **P<0.01, ***P<0.001 and ****P<0.0001. n.s. indicates no significance.

## Acknowledgements

We thank Eric Wieschaus (*His2Av-GFP*), Romain Levayer (*miniCic::mNeonGreen*), and Mike Levine (*MCP::mCherry*) for the kind gift of flies used in this study. We thank Trudi Schüpbach, Mathieu Coppey, and all members of the Toettcher and Shvartsman labs for helpful discussions. Confocal imaging was done in the Princeton Confocal Imaging Facility, a Nikon Center of Excellence, with assistance from Gary Laevsky and Sha Wang. Stocks obtained from the Bloomington Drosophila Stock Center (NIH P40OD018537) were used in this study. We thank QG Bu for assistance with stock maintenance. This work was supported by the National Institutes of Health grants F32GM148016 (E.K.H.), R01HD085870 (S.Y.S.), R01GM141843 (S.Y.S.), R01GM134204 (S.Y.S.), and U01DK127429 (J.E.T.). Research reported in this publication was also supported by the National Center for Advancing Translational Sciences (NCATS), a component of the National Institutes of Health (NIH) under award number TL1TR003019 (Kim-Yip).

## Author Contributions

E.K.H., P.E.F., and J.E.T. designed the study. E.K.H. performed experiments. P.E.F. cloned constructs. R.P.K-Y. performed experiments in egg chambers. A.S. performed experiments in trachea. E.K.H. and H.R.O. analyzed the data. E.P., S.Y.S., and J.E.T. supervised the study. E.K.H and J.E.T. wrote the manuscript with feedback from all authors.

