## Supplemental Materials for "*In vivo* measurements of receptor tyrosine kinase activity reveal feedback regulation of a developmental gradient"

Supplemental Figures S1-S4

Supplemental Figure 1 (Related to Figure 1)
Supplemental Figure 2 (Related to Figure 2)
Supplemental Figure 3 (Related to Figure 3)
Supplemental Figure 4 (Related to Figure 4)

Supplemental Tables S1-S2

Supplemental Table S1 (Related to Material and Methods)
Supplemental Table S2 (Related to Material and Methods)

Legends for Supplemental Tables S1-S2

Supplemental Movie S1 (Related to Figure 3)
Supplemental Movie S2 (Related to Figure 4)

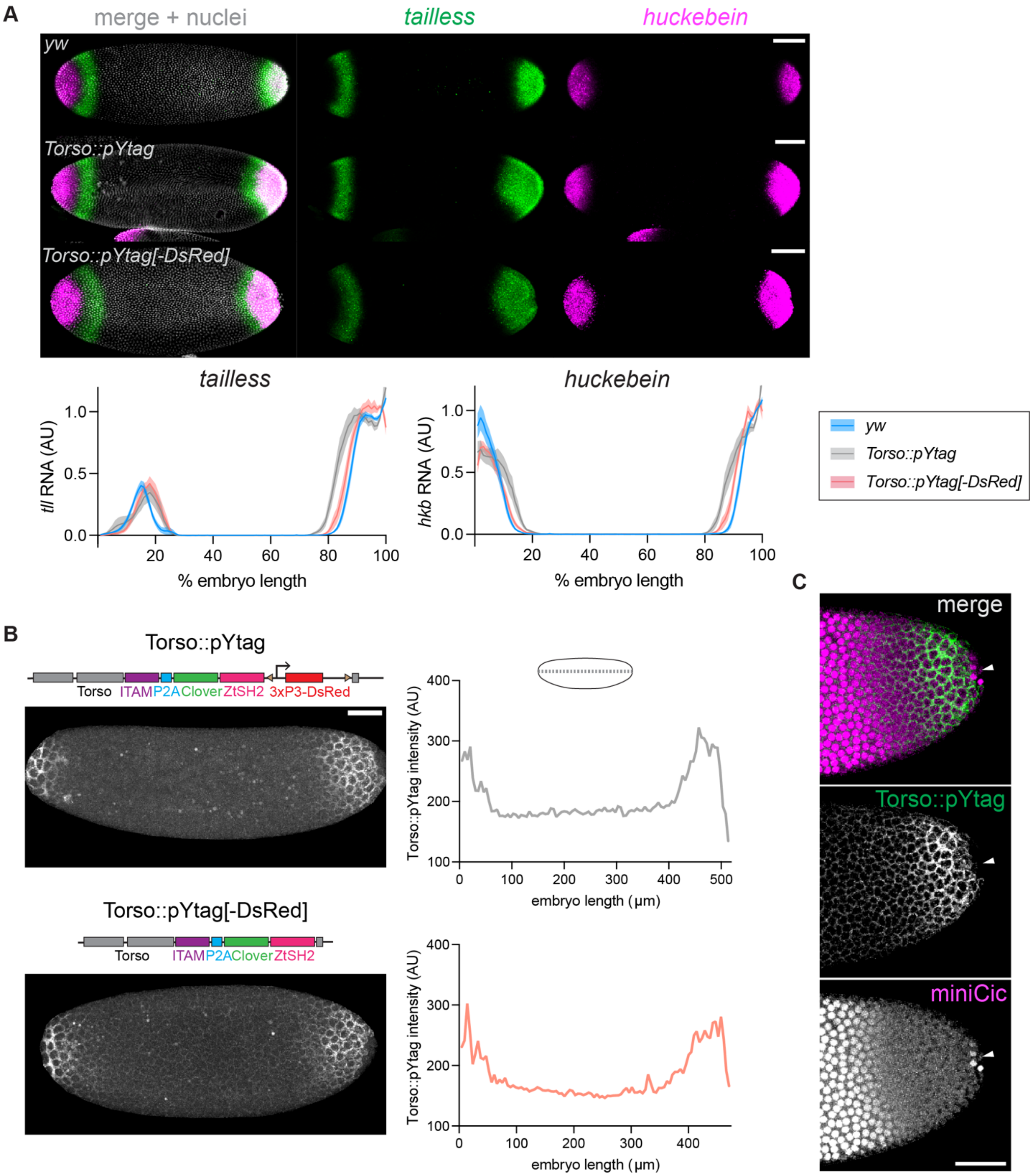


**Figure S1: Torso::pYtag detects Torso activity without significantly disrupting endogenous terminal patterning.** (A) To assess terminal patterning in wild-type (*yw*), *Torso::pYtag*, and *Torso::pYtag[-DsRed]* embryos, NC14 embryos were stained for *tll* and *hkb* mRNA using HCR FISH. Images show representative images and graphs show quantification across the anterior-posterior axis. The boundaries of *tll* and *hkb* expression, which are set by the ERK gradient, shift towards the center of the embryo in *Torso::pYtag* embryos but are largely restored to their wild-type positions when the 3xP3-DsRed marker is removed and only the pYtag is present. This result suggests that pYtag insertion influences terminal patterning but this effect is largely due to the 3xP3-DsRed and not the pYtag itself. Lines are mean ± s.e.m. n = 20 (*yw*), 13 (*pYtag*), 15 (*pYtag[-DsRed]*). Scale bar: 50 μm. (B) Comparison of Torso activity in *Torso::pYtag* and *Torso::pYtag[-DsRed]* embryos confirms that both genotypes had similar intensities and spatial patterns of Torso activity at the poles. Images show NC12 embryos. Bright foci in the center of the embryo are autofluorescent yolk granules. Graphs show the Clover-ZtSH2 intensity across the anterior-posterior axis for the pictured embryo of each genotype. Scale bar: 50 μm. (C) Repression of Torso activity in the pole cells is detectable by Torso::pYtag. Image shows the posterior pole of a NC13 embryos. Arrowhead denotes pole cells which have nuclear miniCic (ERK OFF) and no Clover-ZtSH2 signal. Scale bar: 50 μm.

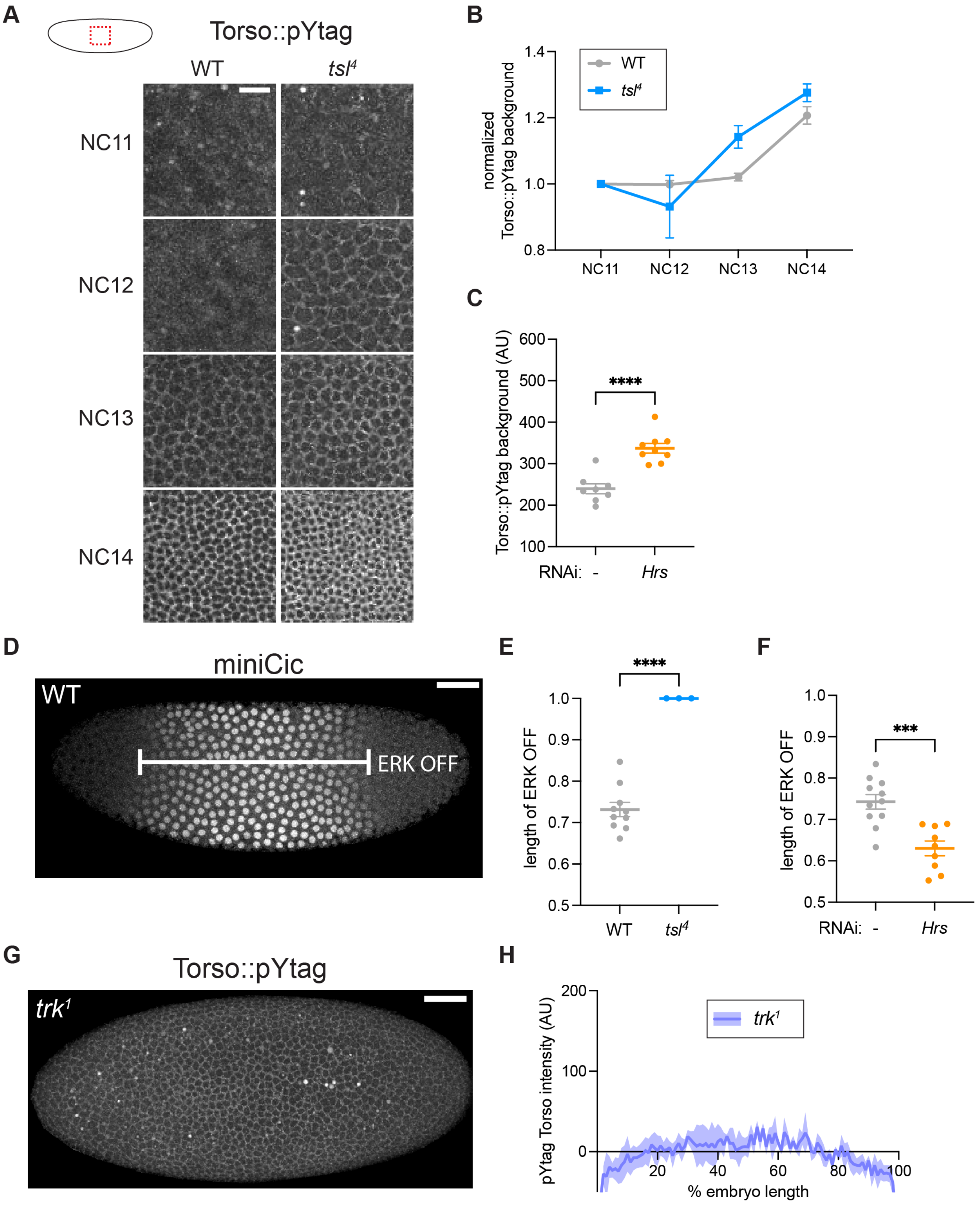


**Figure S2: Validation of genetic perturbations to Torso activity.** (A) Low-level localization of Clover-ZtSH2 to membranes in the center of the embryo where Torso is inactive, shown for WT and *tsl*$^{4}$ embryos in NC11-14. Scale bar: 20 μm. (B) Comparison of background Torso::pYtag intensity in WT and *tsl*$^{4}$ embryos across NC11-14. Background intensities are normalized to the

NC11 value. Mean ± s.e.m. from n = 9, 4 embryos. (C) Comparison of background Torso::pYtag intensity in control and *Hrs* RNAi NC13 embryos shows that *Hrs* RNAi embryos have significantly higher Torso::pYtag background. Mean ± s.e.m., n = 8, 9 embryos. Unpaired t test. (D) To validate genetic perturbations to Torso, we sought a simple measure to compare ERK activity between embryos and conditions. The proportion of embryo length in which miniCic is nuclear reveals the "length of ERK OFF". The image shows an example of the "ERK OFF" portion of a wild-type NC12 embryo. Scale bar: 50 µm. (E) Comparison of ERK OFF embryo length in WT and *tsl*$^{4}$ NC14 embryos. All *tsl*$^{4}$ mutant embryos show nuclear miniCic throughout the entire embryo (see **Figure 2C**), indicating no active ERK signaling. Mean ± s.e.m., n = 10, 3 embryos. Unpaired t test. (F) Comparison of ERK OFF embryo length in control and *Hrs* RNAi NC14 embryos. The *Hrs* RNAi phenotype is variable, but there is a significant decrease in the ERK OFF length compared to controls, indicating higher ERK signaling. Mean ± s.e.m., n = 11, 9 embryos. Unpaired t test. (G) Torso::pYtag in a *trk*$^{1}$ NC13 embryo shows no active Torso at the poles. Like in the *tsl*$^{4}$ mutant, there is low level membrane localization of Torso::pYtag throughout the embryo. Scale bar: 50 µm. (H) Quantification of Torso::pYtag across the anterior-posterior axis of NC13 *trk*$^{1}$ mutant embryos. Mean ± s.e.m. from n = 3 embryos.

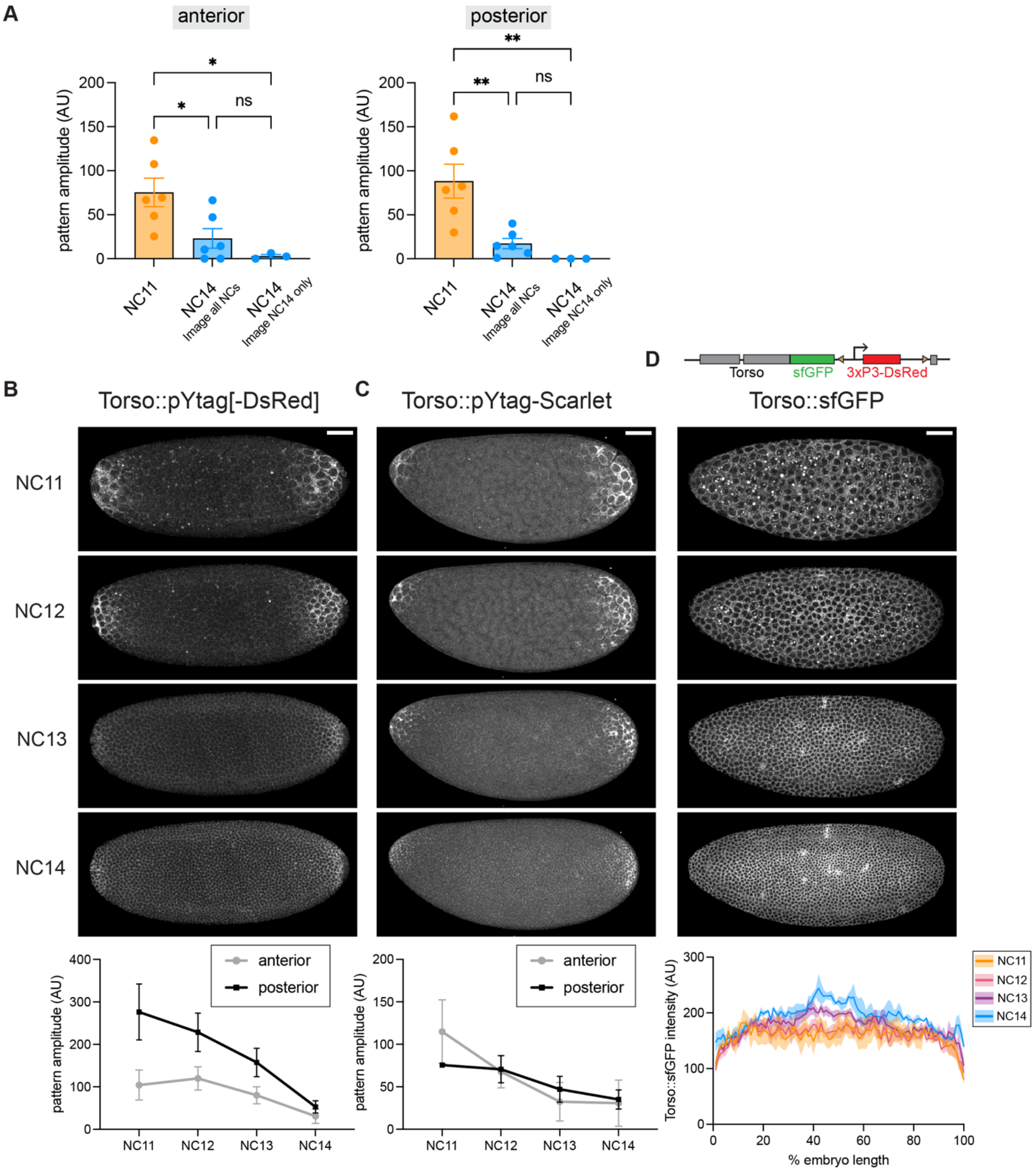


**Figure S3: Validating the decrease in Torso activity over time.** (A) To show that the decrease in Torso::pYtag intensity is not due to bleaching, NC14 embryos that were only imaged in NC14 were compared to embryos that had been imaged in NC11-14. There was no difference in these conditions, and both were significantly decreased compared to Torso::pYtag in NC11. RNAi control embryos were used for this experiment. Mean ± s.e.m. Significance from one-way ANOVA with Tukey's test. n = 3-6 embryos. (B-C) Images show a (B) Torso::pYtag[-DsRed] and (C) Torso::pYtag-Scarlet embryo over NC11-14. In both embryos, Torso::pYtag pattern amplitude decreases over time in the anterior and the posterior. Mean ± s.e.m. n = 4 Torso::pYtag[-DsRed] and n = 7 Torso::pYtag-Scarlet embryos. (D) To visualize total Torso levels, we endogenously tagged the C-terminus of Torso with superfolder GFP (Torso::sfGFP). Images show a Torso::sfGFP embryo over NC11-14. Quantification of Torso::sfGFP across the anterior-posterior axis shows total Torso is slightly increasing over time. The drop-off in signal at the poles is due to the curvature of the embryo and is also observed for Torso::pYtag in $tsl^4$ embryos which do not have signal enrichment at the poles (see **Figure 2D**). All scale bars: 50 µm.

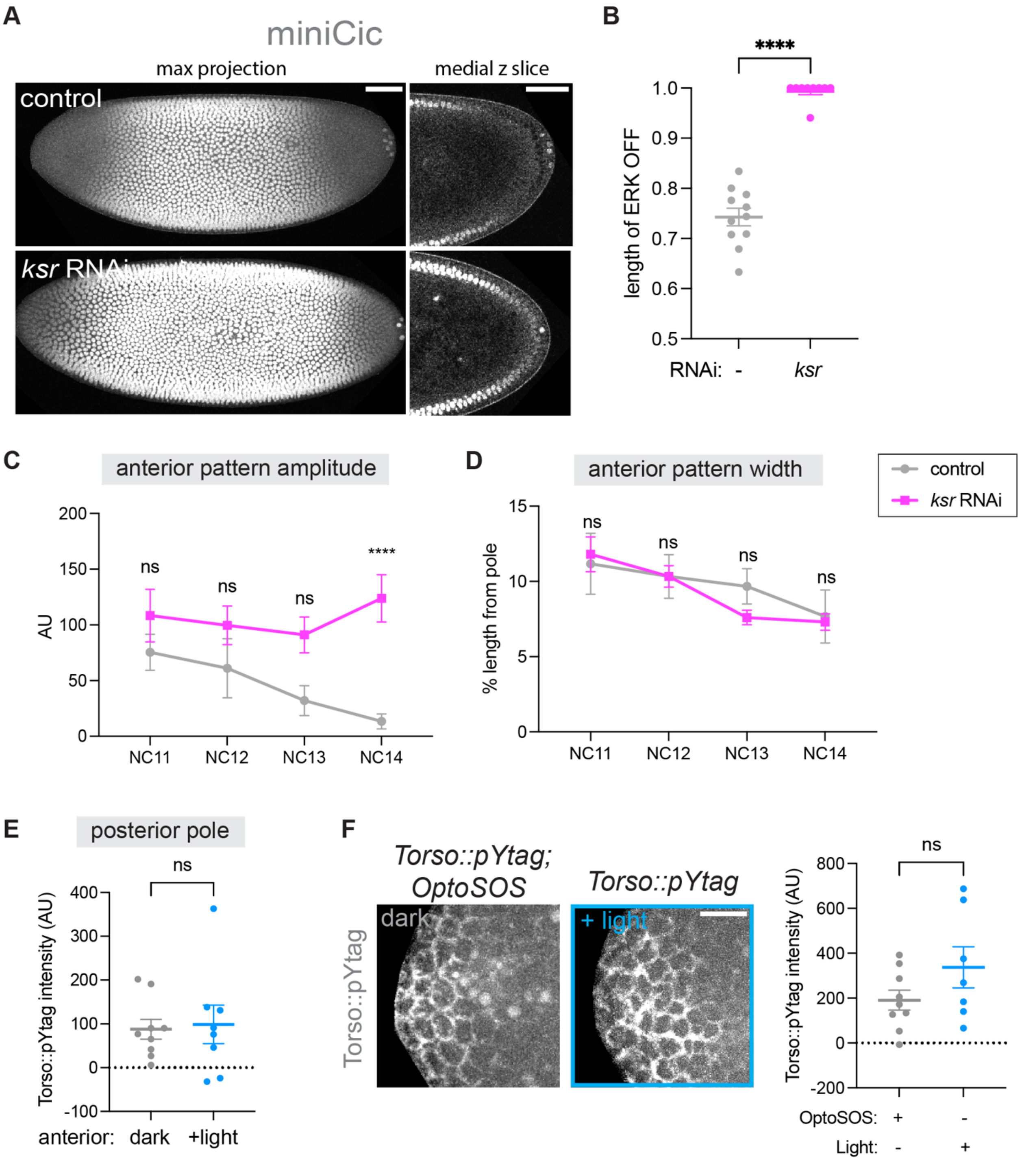


**Figure S4: Validation of manipulations to assess negative feedback.** (A) To validate that the *ksr* RNAi was effective in inhibiting ERK activity, we assessed miniCic activity in NC14 control and *ksr* RNAi embryos. While control embryos have cytoplasmic miniCic at the poles indicating that ERK is active, *ksr* RNAi embryos have nuclear miniCic throughout the embryo. We note that the nuclear intensity at the poles of *ksr* RNAi is lower than in the center of the embryo, suggesting there is a low level of residual ERK activity. However, the medial z slice shows that the signal is nuclear all the way to the pole, revealing that this residual ERK activity is very low. (B)

Quantification of the ERK OFF embryo length in control and *ksr* RNAi NC14 embryos. Mean ± s.e.m. n = 11 (control), 9 (*ksr*) embryos. Significance by unpaired t test. (C) Anterior pattern amplitude over time for the same embryos as in (**Figure 4D**). The anterior pole shows a similar increase in Torso::pYtag activity in *ksr* RNAi embryos over time. Mean ± s.e.m. Significance by 2-way ANOVA with multiple comparisons test. (D) Anterior pattern width over time for the same embryos as in (**Figure 4E**). Mean ± s.e.m. Significance by 2-way ANOVA with multiple comparisons test. (E) Posterior Torso::pYtag intensity for the same embryos as in (**Figure 4H**). The posterior was unilluminated in all embryos and thus shows no difference between conditions. Mean ± s.e.m. n = 9. Unpaired t test. (F) To confirm that optogenetic stimulation with blue light did not bleach the Torso::pYtag, we illuminated Torso::pYtag embryos that did not express OptoSOS with the same illumination regime. There was no significant difference in anterior Torso::pYtag intensity between illuminated Torso::pYtag embryos and unilluminated Torso::pYtag; OptoSOS embryos. Mean ± s.e.m. n = 9 (unilluminated), 7 (illuminated). Unpaired t test.

**Table S1.** Primers used for CRISPR/Cas9 and genotyping

| Primer Name | Sequence |
|---|---|
| Torso 5' Homology F | CAAGAAGCGAATCTTTGAGAACAAGGAATACTTTGATTGCCTCGACTCATCGG |
| Torso 5' Homology R | ATTCAAAGGTTCTAGGTATAGCTCTTCCTCGCATGGCACTTGC |
| Torso 3' Homology F | TAGTCATTGCTTCAAGATTATAATGAACGAGTGCAATACATTCTAAATTCGAGTTCC |
| Torso 3' Homology R | CTAAGGGCGCTCAGGAGCTTTGGAATGGACACAACATCG |
| Torso Guide F | cttcgAGCAATGACTATTAATTCAA |
| Torso Guide R | aaacTTGAATTAATAGTCATTGCTc |
| Torso Genotyping F | AATAACCAATGCAGCCGACAACAAGGGCTATGGCCTGG |
| Torso Genotyping R | GGACTCTTTGGTTCCGTCACTTCGGAGTAGAGATCAGATATACTTCTCACC |
| EGFR 5' Homology F | TGAGTACAAGGCTGCTGGCGGCAAGATGCCCATCAAGTGG |
| EGFR 5' Homology R | CACCCTCGTCTCCGTGTTGCGGTTTCGATGCAGTGG |
| EGFR 3' Homology F | GCTCCAGTCGAGTAGGAGCAATTGCCAATGAAGAAGGAGAATCTTGCC |
| EGFR 3' Homology R | TTCTTGGCGGGCACCAACCGGTTATCAAGCC |
| EGFR PAM SDM F | AGACGAGaGTGGGCTCtGGtTCTCCACCTCCC |
| EGFR PAM SDM 5 | AGCCCACtCTCGTCTCCGTGTTGCGGTTTCG |
| EGFR Guide F | cttcGAAACCGCAACACGGAGACG |
| EGFR Guide R | aaacCGTCTCCGTGTTGCGGTTTC |
| EGFR Genotyping F | AAGATCACCGACTTTGGGCTGGCCAAGTTGC |
| EGFR Genotyping R | ACGAAATACAGTTTGCGACCACGCCCTCTATAGAACAACC |
| Btl 5' Homology F | AAGATGGTCAAGGAGGAGCACACGGATACGGACATGG |
| Btl 5' Homology R | AGGTGTACTGATATCTCAGTGGAGACGTTTCCCGGAATGTTTCGGTGTCGGAGCC |
| Btl 3' Homology F | TTCGTAGTATAAGGAGACCAAAAAGAATTCCAACGAGTCAATCAGATCCCATCGAAGC |
| Btl 3' Homology R | GCTGTCAGGATCATCGTTAAGTTGGCTCCCCATTGTAATGAGTTCCTCG |
| Btl Guide F | cttcGAAACCGCAACACGGAGACG |
| Btl Guide R | aaacCGTCTCCGTGTTGCGGTTTC |
| Btl Genotyping F | AGCAGCTTAGCTTGGGCTCCATTTTGGGTGAGG |
| Btl Genotyping R | CGATAGATTCCCAACAAATTCAGGGGCATTTCCTCATCGAGCTG |

**Table S2**. Fly genotypes used

| Figure | Maternal genotype |
|---|---|
| 1D, 1E, S1A-C, 2A, 2C, 2D, S2A, S2B, S2D, S2E, 3B-E, S4F | Torso::pYtag, miniCic::mCherry |
| 1D, 1E | EGFR::pYtag, miniCic::mCherry |
| 1D, 1E | Zygotic genotype: btl-Gal4, UAS-CD4::mIFP / CyO ; btl::pYtag |
| S1A, 3B-E | yw |
| S1A, S1B, S3B | Torso::pYtag[-DsRed]; CAAX-mCherry |
| 2A, 2C, 2D, S2A, S2B, S2E | Torso::pYtag, miniCic::mCherry / Torso::pYtag ; $tsl^4$ / 15, $tsl^4$ |
| 2B | Torso::pYtag[-ITAM], miniCic::mCherry |
| 2E, 2F, S2C, S2F, S3A, 4B-E, S4A-D | Torso::pYtag, 67 / Torso::pYtag, miniCic::mCherry ; 15 / + |
| 2E, 2F, S2C, S2F | Torso::pYtag, 67 / Torso::pYtag, miniCic::mCherry ; 15 / UAS-Hrs RNAi |
| S2G, S2H | $trk^1$, Torso::pYtag |
| 3B-E | miniCic::mCherry / + ; His2Av-GFP / + |
| S3C | Torso::pYtag-Scarlet, miniCic::mNeonGreen |
| S3D | Torso::sfGFP / CyO |
| 4B-E, S4A-D | Torso::pYtag, 67 / Torso::pYtag, miniCic::mCherry ; 15 / UAS-ksr RNAi |
| 4G, 4H, S4E, S4F | Torso::pYtag, 67 / Torso::pYtag ; 15 / UAS-optoSOS |
| 5A, 5B | Torso::pYtag, MCP::mCherry / Torso::pYtag x Sp/CyO ; tll-MS2 males |

### Movie Captions

**Movie S1: Wild-type Torso::pYtag dynamics in embryonic posterior**
Movie shows the posterior of a wild-type embryo expressing Torso::pYtag imaged at 3 minute intervals from NC11 through NC14. Timer (hh:mm) shows time since imaging began in NC11. Each frame is labeled with the corresponding nuclear cycle (NC). Scale bar is 50 μm.

**Movie S2: *ksr* RNAi Torso::pYtag dynamics in embryonic posterior**
Movie shows the posterior of a *ksr* RNAi embryo expressing Torso::pYtag imaged at 3 minute intervals from NC11 through NC14. Timer (hh:mm) shows time since imaging began in NC11. Each frame is labeled with the corresponding nuclear cycle (NC). Scale bar is 50 μm.
